# Defects in DNA double-strand break repair re-sensitise antibiotic-resistant *Escherichia coli* to multiple bactericidal antibiotics

**DOI:** 10.1101/2022.01.24.477632

**Authors:** Sarah A. Revitt-Mills, Elizabeth K. Wright, Madaline Vereker, Callum O’Flaherty, Fairley McPherson, Catherine Dawson, Antoine M. van Oijen, Andrew Robinson

**Affiliations:** Molecular Horizons Institute and School of Chemistry and Molecular Bioscience, University of Wollongong, Wollongong, Australia; Illawarra Health and Medical Research Institute, Wollongong, Australia

## Abstract

Antibiotic resistance is becoming increasingly prevalent amongst bacterial pathogens and there is an urgent need to develop new types of antibiotics with novel modes of action. One promising strategy is to develop resistance-breaker compounds, which inhibit resistance mechanisms and thus re-sensitise bacteria to existing antibiotics. In the current study, we identify bacterial DNA double-strand break repair as a promising target for the development of resistance-breaking co-therapies. We examined genetic variants of *Escherichia coli* that combined antibiotic-resistance determinants with DNA repair defects. We observed that defects in the double-strand break repair pathway led to significant re-sensitisation towards five bactericidal antibiotics representing different functional classes. Effects ranged from partial to full re-sensitisation. For ciprofloxacin and nitrofurantoin, sensitisation manifested as a reduction in the minimum inhibitory concentration. For kanamycin and trimethoprim, sensitivity manifested through increased rates of killing at high antibiotic concentrations. For ampicillin, repair defects dramatically reduced antibiotic tolerance. Ciprofloxacin, nitrofurantoin, and trimethoprim induce the pro-mutagenic SOS response. Disruption of double-strand break repair strongly dampened the induction of SOS by these antibiotics. Our findings suggest that if break-repair inhibitors can be developed they could re-sensitise antibiotic-resistant bacteria to multiple classes of existing antibiotics and may supress the development of de novo antibiotic-resistance mutations.

## Introduction

The emergence of antimicrobial resistance (AMR) poses a significant global health threat, with once trivial bacterial infections becoming increasingly difficult to treat ^1^. AMR has rendered several current antibiotics effectively obsolete, severely limiting infection treatment options ^2, 3^. There is significant interest in developing combinational drugs that can extend the clinical lifetimes of current therapeutics ^4, 5^. One possibility is the development of ‘resistance breaking’ compounds that increase sensitivity to current antimicrobial therapies ^6, 7^.

There is growing evidence that treatment with certain antibiotics can elevate mutation rates in bacteria, potentially increasing the likelihood that antibiotic resistance mutations will appear in bacterial populations ^8–11^. For any new therapy it would be desirable to limit the possibility of mutation by i) narrowing the mutant selection window ^12^ and ii) by suppressing mutagenesis ^13^. The current study identifies bacterial DNA double strand break repair (DSBR) as a promising target for the development of such therapies.

Several commonly used bactericidal antibiotics have been shown to damage bacterial DNA either as a direct consequence of their primary mode of action, or through secondary effects ^14^. Many forms of DNA damage are lethal to bacteria if left unrepaired ^15^. Bacteria have sophisticated systems to repair DNA damage and the action of these repair pathways effectively offsets killing by DNA-damaging antibiotics ^16^. Recent studies have demonstrated that inactivation of bacterial DNA repair pathways can sensitise bacterial cells to multiple antibiotics. In *E. coli*, inactivation of *recA,* a key contributor to DNA repair via homologous recombination, has been shown to reduce minimum inhibitory concentrations (MICs) against the antibiotics ceftazidime (β-lactam; cephalosporin), fosfomycin (phosphonic antibiotic), ciprofloxacin (quinolone), trimethoprim (dihydrofolate synthesis inhibitor), and colistin (polymixin) ^17^. Promisingly, deletion of *recA* also re-sensitised a ciprofloxacin-resistant strain of *E. coli* to clinically approachable levels of ciprofloxacin ^18^. The *recA* gene is required for the repair of both double-stranded DNA breaks and single-stranded DNA gaps ^19^. It is unclear whether these antibiotic-sensitising effects stem from defects in the DSBR or single-strand gap repair (SSGR) pathways. It is also unclear whether these re-sensitisation effects extend to other classes of bactericidal antibiotics. In this study, we aim to address these shortfalls by measuring MICs, examining time-kill kinetics, and determining antibiotic tolerance phenotypes for *E. coli* strains defective in DSBR and SSGR. Five antibiotics documented as having bactericidal effects were examined: ciprofloxacin ^20^, nitrofurantoin ^21^, kanamycin ^22^, trimethoprim ^23^ and ampicillin ^24^.

In *E. coli* double-strand DNA breaks are primarily repaired through homologous recombination via the RecA protein and RecBCD pathway ^25^. Single-strand gaps are predominantly repaired by RecF, RecO and RecR proteins through their aiding in RecA-mediated homologous recombination ^26^. A third pathway that is utilised under DNA damage conditions is nucleotide pool sanitation (NPS). The NPS pathway removes oxidized nucleotides from the resource pool and thus prevents insertion of aberrant bases during DNA synthesis ^27^. In the absence of one particular NPS enzyme, MutT, insertion of the aberrant base 8-oxo-dGTP into the DNA triggers a form of maladaptive DNA repair that can kill bacterial cells ^23^. We examined the effects of disrupting MutT alongside the DSBR and SSGR pathways in this work.

DNA damage also induces a mutation-promoting stress-response mechanism called the SOS response ^28^. In some circumstances, induction of the SOS response has been observed to increase the frequency of antibiotic-resistance mutations that appear in bacterial populations^13, 29^. Among the ∼40 genes induced during SOS are genes that encode error-prone DNA polymerases known to cause an array of mutations ^30^. SOS is induced by RecA* nucleoprotein filaments that form in response to DNA damage ^31^. Through disruption of *recA* and other DNA-repair genes there is potential to attenuate SOS mutagenesis. Enhancing killing and decreasing mutation supply through inhibition of bacterial DNA repair pathways represents a global approach towards re-sensitisation of antibiotic resistant bacteria. In this study, we measure induction of the SOS response by each of the five antibiotics and examine the effects of disrupting DSBR and SSGR on SOS induction.

## Results

### Disrupting DSBR re-sensitises ciprofloxacin-resistant and nitrofurantoin-resistant *E. coli*

It is widely accepted that ciprofloxacin and other fluoroquinolones induce DNA damage in bacteria ^20, 21^. Ciprofloxacin targets the essential bacterial enzymes DNA gyrase and topoisomerase IV, stabilising a protein-bridged DNA double-strand break intermediate that is formed during supercoiling and decatenation reactions ^20, 32–34^. Cells treated with ciprofloxacin are known to accumulate DNA double-strand breaks ^20, 32^. The mechanism(s) underlying the formation of ciprofloxacin-induced breaks remain under investigation ^33, 35^. Resistance to ciprofloxacin commonly develops through the acquisition of mutations in the genes encoding DNA gyrase and topoisomerase IV. Ciprofloxacin has reduced affinity for these mutant forms of DNA gyrase and topoisomerase IV ^36, 37^. In this study we made use of a ciprofloxacin-resistant (Cip^R^) derivative of *E. coli,* CH5741 ^38^. This strain has clinically relevant point mutations in the genes of the quinolone targets: DNA gyrase (*gyrA;* [S83L, D87N]) and topoisomerase IV (*parC;* [S80I]). Introduction of these point mutations increases ciprofloxacin MIC 1000-fold in comparison to the sensitive background ^38^. It is assumed that double-strand breaks are still formed in this background, but due to reduced target affinity, far higher concentrations of ciprofloxacin would be required.

Reasoning that bacterial DNA repair might offset the killing effects of ciprofloxacin in both sensitive and resistant backgrounds, we examined the sensitivities of *E. coli* strains that combined defects in the DSBR, SSGR and NPS repair pathways with ciprofloxacin-resistance mutations. We determined the MIC for each strain using Liofilchem® MTS™ (MIC Test Strips). Cells deficient in RecA or RecB were hypersensitive to ciprofloxacin in comparison to the wild-type (ciprofloxacin-sensitive) background (Figure 1a). This phenotype was rescued upon complementation with *recA* and *recB* in trans (Figure 1b), confirming the involvement of both RecA and the DSBR pathway in ciprofloxacin sensitivity. Cells lacking the SSGR protein RecO initially appeared to be more sensitive to ciprofloxacin than wild-type cells, but this was found not to be statistically significant. Deletion of other genes whose products are involved in single-stranded DNA repair (*recF,* and *recR)* or nucleotide sanitation (*mutT*) had no significant effect on ciprofloxacin MIC. These findings indicate that DSBR is required for the repair of DNA damage following exposure to ciprofloxacin, in agreement with previous studies ^32, 39^.

**Figure 1:**
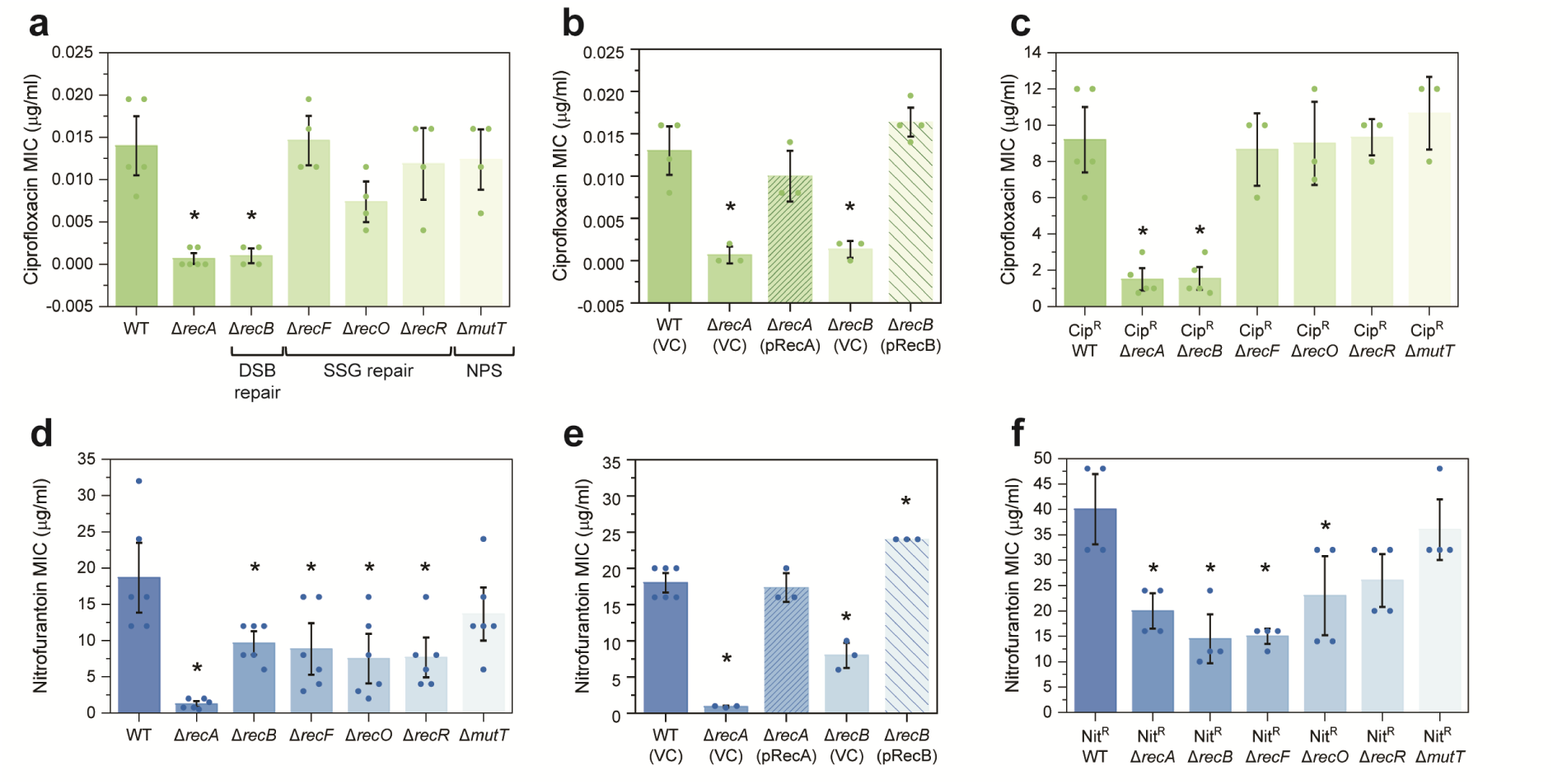
Cells deficient in double-strand break repair have an increased sensitivity to ciprofloxacin and nitrofurantoin. **a.** Ciprofloxacin MIC values obtained for isogenic *E. coli* strains MG1655 (WT; wild-type), Δ*recA*::Kan^R^ (HH020), Δ*recB*::Kan^R^ (EAW102), Δ*recF*::Kan^R^ (EAW629), Δ*recO*::Kan^R^ (EAW114), Δ*recR*::Kan^R^ (EAW669) and Δ*mutT*::Kan^R^ (EAW999). The means and standard errors of the mean are shown, based on results from at least four biological replicates. Statistical analysis was carried out using two sample students *t*-tests. An asterisk denotes statistical significance (*p* < 0.05) compared to wild-type (MG1655). **b.** Ciprofloxacin MIC values obtained for the *E. coli* strains wild-type (MG1655) with empty vector (VC; vector control), Δ*recA* and Δ*recB* mutants with empty vector and complemented derivatives (pRecA and pRecB, respectively). The means and standard errors of the mean are shown based on results from at least three biological replicates. Statistical analysis was carried out using two sample students *t*-tests. An asterisk denotes statistical significance (*p* < 0.05) compared to wild-type with empty vector, WT (VC). **c.** Ciprofloxacin MIC values obtained for isogenic ciprofloxacin-resistant (Cip^R^) DNA repair deficient *E. coli* strains Cip^R^ (CH5741), Cip^R^ Δ*recA* (FM002), Cip^R^ Δ*recB* (FM001), Cip^R^ Δ*recF* (FM003), Cip^R^ Δ*recO* (FM004), Cip^R^ Δ*recR* (FM005) and Cip^R^ Δ*mutT* (MV001). The means and standard errors of the mean are shown, based on results from at least three biological replicates. Statistical analysis was carried out using two-sample student’s *t*-tests. An asterisk denotes statistical significance (*p* < 0.05) compared to Cip^R^. **d**. Nitrofurantoin MIC values obtained for isogenic *E. coli* strains. The means and standard errors of the mean are shown, based on results from at least six biological replicates. Statistical analysis was carried out using student’s *t*-tests. An asterisk denotes statistical significance (*p* < 0.05) compared to wild-type (MG1655). **e**. Nitrofurantoin MIC values obtained for the *E. coli* strains wild-type (MG1655) with empty vector (VC; vector control), Δ*recA* and Δ*recB* mutants with empty vector and complemented derivatives (pRecA and pRecB, respectively). The means and standard errors of the mean are shown based on results from at least three biological replicates. Statistical analysis was carried out using student’s *t*-tests. An asterisk denotes statistical significance (*p* < 0.05) compared to wild-type with empty vector, WT (VC). **f.** Nitrofurantoin MIC values obtained for isogenic nitrofurantoin-resistant (Nit^R^) DNA repair deficient *E. coli* strains Nit^R^ (EKW046), Nit^R^ Δ*recA* (EKW047), Nit^R^ Δ*recB* (EKW048), Nit^R^ Δ*recF* (EKW049), Nit^R^ Δ*recO* (EKW050), Nit^R^ Δ*recR* (EKW051) and Nit^R^ Δ*mutT* (EKW052). Statistical analysis was carried out using student’s *t*-tests. The means and standard errors of the mean are shown, based on results from at least three biological replicates. An asterisk denotes statistical significance (*p* < 0.05) compared to Nit^R^.

To determine if these findings translated to an antibiotic-resistant background, the sensitisation effect of disrupting DNA repair genes was examined using a ciprofloxacin resistant (Cip^R^) derivative of *E. coli,* CH5741 ^38^. Deletion of *recA* and *recB* led to significant re-sensitisation towards ciprofloxacin, reducing the respective MICs 7-fold and 6-fold in comparison to the ciprofloxacin-resistant parental strain (Figure 1c). Disruption of SSGR and NPS did not alter MIC in the Cip^R^ background. These results support our assumption that double-strand breaks are formed in the Cip^R^ background and indicate that disruption of DSBR reduces resistance in *E. coli* that have already acquired Cip^R^ mutations.

Nitrofurantoin is seldom used as an antibiotic as it has toxic side effects, however its low propensity for resistance has rekindled interest in its use ^40^. The mechanism of action for nitrofurantoin is poorly understood. Studies suggest two mechanisms: i) inhibition of ribosomes, and consequently, protein synthesis ^21^; ii) direct damage to DNA ^41^. Nitrofurantoin is a prodrug. Conversion from the prodrug to active drug form requires nitrofurantoin to be processed intracellularly by the bacterial nitroreductases NfsA and NfsB ^42^. Resistance to nitrofurantoin is associated with loss-of-function mutations in these two nitroreductases, which results in the drug remaining in the inactive prodrug state ^43^. In the current study, nitrofurantoin-resistant (Nit^R^) strains were constructed through deletion of *nfsA,* which encodes the nitroreductase NfsA. At the outset of the study, it was not known whether this mutation would eliminate nitrofurantoin-induced DNA damage or not.

In the nitrofurantoin-sensitive background, both Δ*recA* and Δ*recB* cells were found to be hypersensitive to nitrofurantoin (Figure 1d). Deletion of *recA* resulted in a 15x reduction in MIC from 18.7±2.9 μg/ml (wild-type) to 1.25±0.3 μg/ml. Cells deficient in DSBR (Δ*recB*) demonstrated an MIC half that of the wild-type strain (9.7±0.8 μg/ml). This effect was then rescued upon complementation (Figure 1e). The deletion of the genes *recF*, *recO* or *recR*, which encode SSGR proteins, also significantly reduced MIC compared to wild-type. Deletion of nucleotide sanitation (Δ*mutT)* had no effect on MIC. In the nitrofurantoin resistant (Nit^R^) background, deletion of *recA, recB* or *recF* led to significant re-sensitisation (Figure 1f). Deletion of *recA* fully re-sensitised Nit^R^ cells to wild-type level (20±2.6 μg/ml). Deletion of *recB* or *recF* was even more sensitising, reducing the MIC to below wild-type levels (14.5±3.7 μg/ml and 15±1.2 μg/ml, respectively). Cells lacking other SSGR proteins, RecO and RecR, in addition to MutT showed no significant re-sensitisation. The potent re-sensitisation effects of *recA*, *recB*, and *recF* mutations strongly suggest that DNA damage still occurs in cells that have developed nitrofurantoin resistance through loss of function mutations in NfsA. Disruption of the DSBR pathway fully re-sensitises NfsA-lacking cells activity towards nitrofurantoin.

### Disrupting DSBR enhances killing by kanamycin and trimethoprim

The antibiotics kanamycin and trimethoprim target essential components of the bacterial cell, namely ribosomes and folate biosynthesis ^14, 44^. Whilst the primary action of these antibiotics do not directly induce DNA damage, there is now growing evidence that treatment of bacterial cells with bactericidal antibiotics results in the overproduction of reactive oxygen species (ROS) ^45^. It has been suggested that treatment with these antibiotics provokes accumulation of ROS leading to DNA damage ^35, 45–47^. We therefore examined whether cells lacking DSBR, SSGR, and NPS are sensitised to kanamycin and trimethoprim.

We first examined if DNA repair deficient strains had altered sensitivity to kanamycin by MIC tests. For most strains, MICs were similar to wild type (Supplementary Figure S1a). Disruption of DSBR, SSGR, or NPS did not lead to reduced MICs; in fact MICs were marginally increased in Δ*recO* (1.9±0.2 µg/ml) and Δ*recR* (1.8±0.2 µg/ml) mutants compared to wild-type (1.2±0.3 µg/ml). We did notice, however, that *recA* and *recB* mutants had significantly larger zones of clearing surrounding the MIC strip (Figure 2a and 2b). Complementation of the *recA* and *recB* mutants, did not alter MIC (Supplementary Figure S1b), but did reduce the area within the zone of inhibition (ZOI) (Figure 2c and Supplementary Figure S1c). We hypothesised that the enlarged ZOI might relate to improved clearance of bacterial cells at high drug concentrations. To test this idea, we used time-kill assays to examine killing of our *E. coli* strains following exposure to 3×, 5× or 10× MIC kanamycin. We observed no significant killing of any strain at 3× or 5× MIC, however at 10× MIC (Figure 2d) there was significant killing of both Δ*recA* and Δ*recB* mutants. Complementation alleviated this sensitivity (Figure 2e). The increased sensitivity of the *recA* and *recB* mutants highlights the importance of RecA and DSBR in mediating survival following kanamycin treatment.

**Figure 2:**
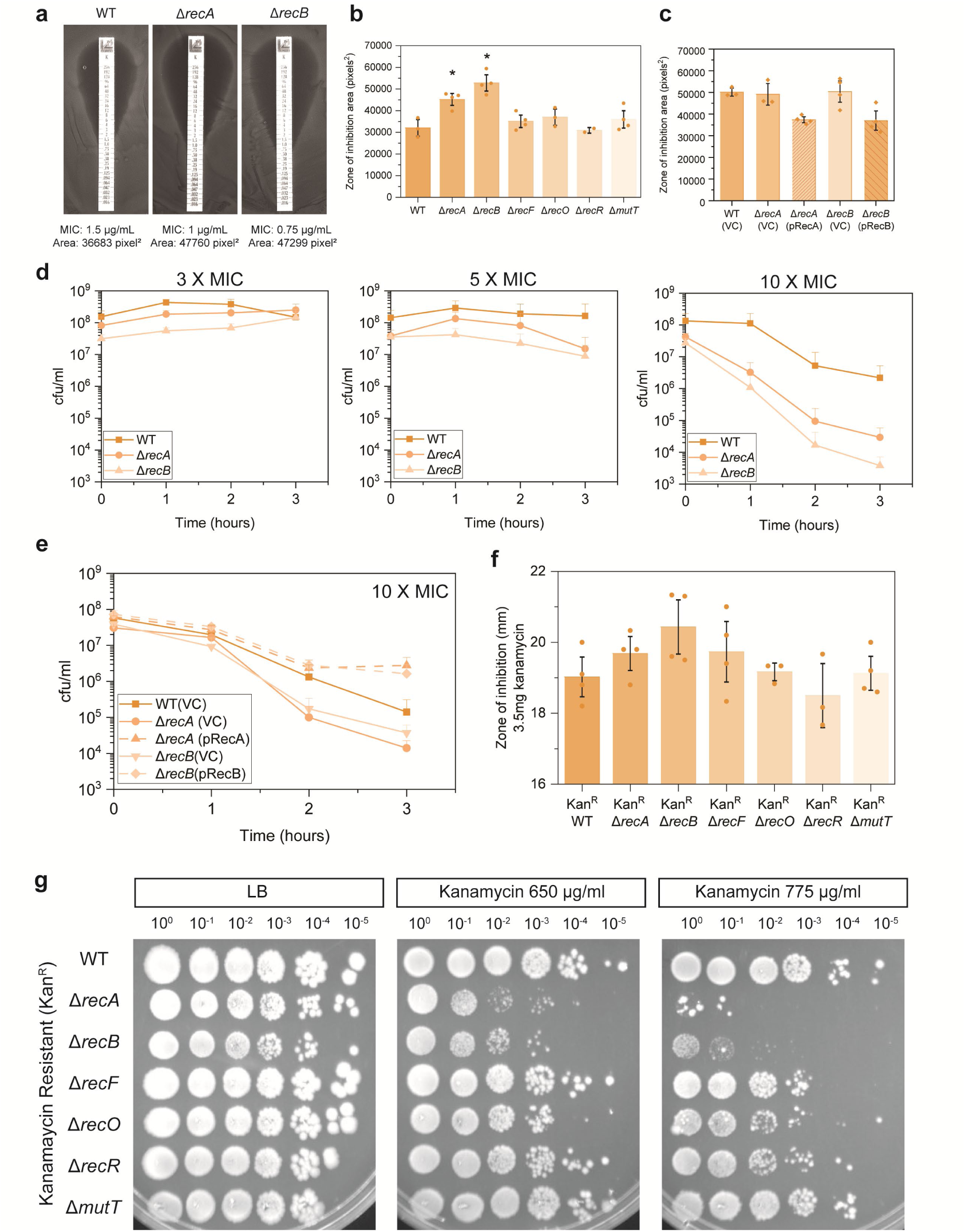
Deletion of double-strand break repair results in moderate sensitivity to the aminoglycoside kanamycin. **a.** Representative images of kanamycin MIC plate assays for wild-type (WT), Δ*recA* and Δ*recB E. coli* strains. MIC’s and measured zone of inhibition areas in pixels^2^ are denoted below the images. **b.** Zone of inhibition area measurements for isogenic *E. coli* strains MG1655 (WT; wild-type), Δ*recA*::FRT (HH021), Δ*recB*::FRT (HG356), Δ*recF*::FRT (MV009), Δ*recO*::FRT (SRM019), Δ*recR*::FRT (SRM020) and Δ*mutT*::FRT (MV005). ZOI areas surrounding MIC strips were measured using ImageJ (Schneider et al., 2012). The means and standard errors of the mean are shown based on results from at least three biological replicates. Statistical analysis was carried out using a student’s *t*-test. An asterisk denotes statistical significance (*p* < 0.05) compared to wild-type (WT; MG1655). **c.** Kanamycin ZOI area values obtained for wild-type (MG1655) with empty vector (VC; vector control), Δ*recA* and Δ*recB* mutants with empty vector and complemented derivatives (pRecA and pRecB, respectively). The means and standard errors of the mean are shown based on results from at least three biological replicates. **d.** Time-kill survival assays for isogenic *E. coli* strains MG1655 (WT; wild-type), Δ*recA*::FRT, Δ*recB*::FRT. Cell viability (cfu/ml) was measured hourly for 3 hours following treatment with 3× MIC (3 µg/ml), 5× MIC (5 µg/ml) or 10× MIC (10 µg/ml) kanamycin. The means and standard errors of the mean are shown based on results from at least three biological replicates. **e.** Complemented time-kill survival assay (10× MIC) for wild-type (MG1655) with empty vector (VC; vector control), Δ*recA* and Δ*recB* mutants with empty vector and complemented derivatives (pRecA and pRecB, respectively). The means and standard errors of the mean are shown based on results from at least three biological replicates. **f.** Zone of inhibition area measurements for kanamycin-resistant WT and DNA repair mutant strains following disk diffusion assays with 3.5 mg kanamycin. Kan^R^ (CD001), Kan^R^ Δ*recA* (CD002), Kan^R^ Δ*recB* (CD003), Kan^R^ Δ*recF* (CD004), Kan^R^ Δ*recO* (SRM026), Kan^R^ Δ*recR* (SRM027) and Kan^R^ Δ*mutT* (CD005). The means and standard errors of the mean are shown based on results from at least three biological replicates. **g**. Spot plate dilution assays of kanamycin-resistant DNA repair deficient strains. Normalised exponential phase cells (OD_600_ 0.2) were diluted to 10^-5^ and spotted onto LB and LB agar supplemented with 650 or 775 µg/ml kanamycin. Images show representative plates from independent triplicate replicates.

We next assessed if this increased sensitivity to kanamycin was also observed in a kanamycin resistant (Kan^R^) background. Kan^R^ derivatives of the DNA repair defective strains were constructed by transformation with the plasmid pUA66, which confers kanamycin resistance through the *aph(3’)-*II gene ^48^. Sensitivity was first assessed via disc diffusion assays with 3.5 mg of kanamycin. No changes in sensitivity to kanamycin were seen in Kan^R^ cells lacking components of SSGR or NPS. The ZOI appeared larger for Kan^R^ *recA* and *recB* mutants, however neither were statistically significant (Figure 2f). As a more sensitive test, viability was assessed using a spot dilution assay. Both the Kan^R^ Δ*recA* and Δ*recB* mutants demonstrated increased sensitivity when plated on 650 µg/ml of kanamycin, with significantly greater sensitivity observed at 775 µg/ml (Figure 2g). No changes in viability were observed for the other strains tested. These findings confirm that both RecA and RecB are required for survival of Kan^R^ cells at high drug-concentrations.

Trimethoprim is a bactericidal drug that disrupts folic acid biosynthesis by inhibiting the enzyme dihydrofolate reductase (DHFR) ^44^. Inhibition of DHFR eventually starves the cell of nucleotides ^49^. Killing by trimethoprim in many ways mirrors the well-studied phenomenon of thymineless death ^50^. As thymineless death is hypothesised to involve the formation of both double-strand breaks and single-strand gaps ^23, 50^, we examined DNA repair deficient strains of *E. coli* for sensitivity to trimethoprim. The Δ*recA* and Δ*recB* mutants showed no significant sensitivity in the MIC assay (Figure 3a). The SSGR mutant Δ*recO* showed sensitivity (0.27±0.04 µg/ml), whereas Δ*recR* demonstrated significant resistance (2.36±0.35 µg/ml) to trimethoprim treatment. We also determined cell viability using a more sensitive spot plate dilution assay (Figure 3b). Increased sensitivity to 0.2 µg/ml trimethoprim was observed for strains lacking RecA or RecB, with the greatest sensitivity observed for the RecO deficient strain. No changes in viability were seen in the other tested mutant strains when compared to wild-type cells.

**Figure 3:**
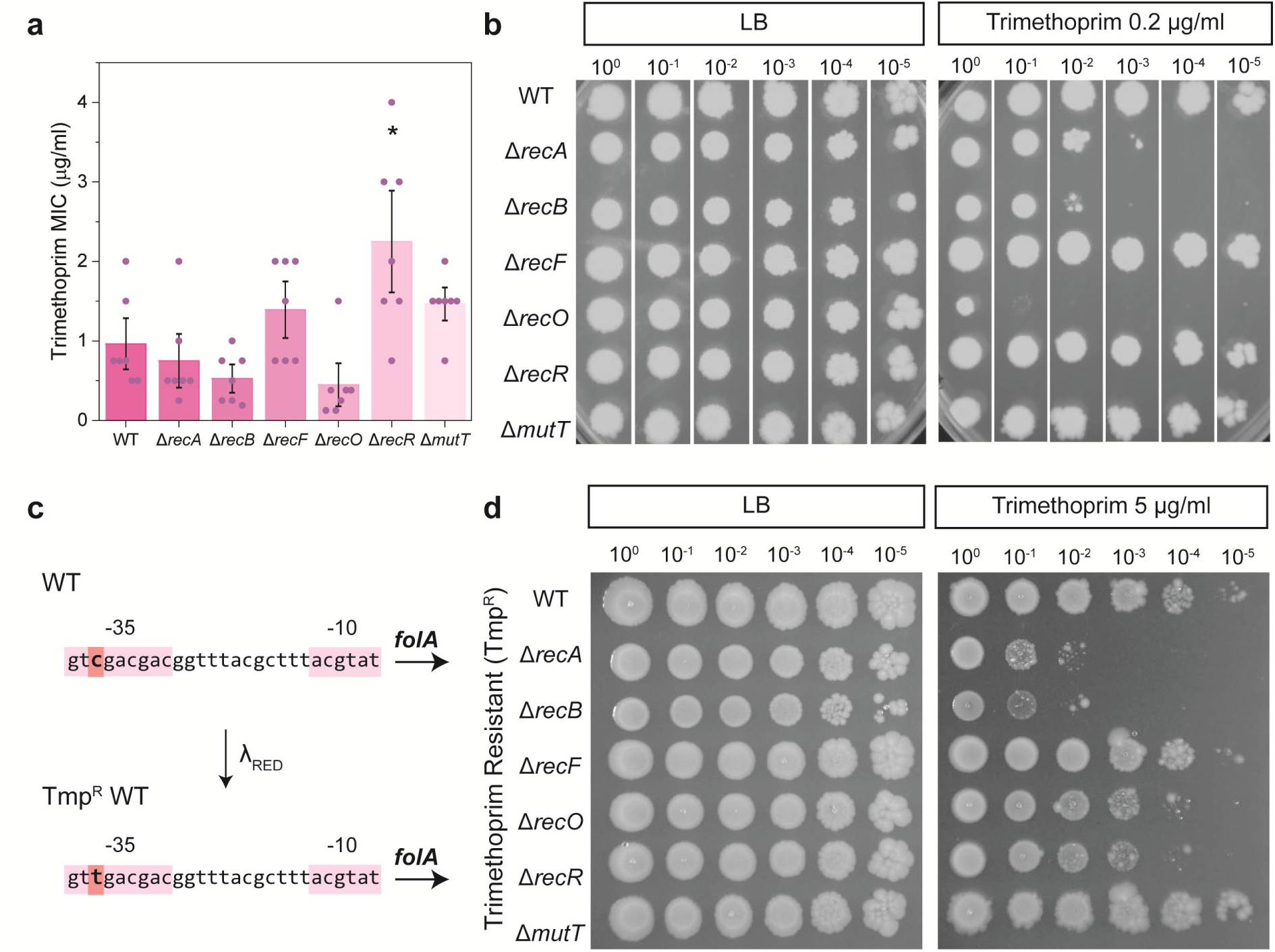
*E. coli* lacking key components of DNA repair pathways demonstrate increased trimethoprim sensitivity. **a.** Trimethoprim MIC values obtained for isogenic *E. coli* strains MG1655 (WT; wild-type), Δ*recA*::Kan^R^ (HH020), Δ*recB*::Kan^R^ (EAW102), Δ*recF*::Kan^R^ (EAW629), Δ*recO*::Kan^R^ (EAW114), Δ*recR*::Kan^R^ (EAW669) and Δ*mutT*::Kan^R^ (EAW999). The means and standard errors of the mean are shown, based on results from at least four biological replicates. Statistical analysis was carried out using a student’s *t*-test. An asterisk denotes statistical significance (*p* < 0.05) compared to wild-type (MG1655). **b.** Spot plate dilution assays of antibiotic-sensitive DNA repair deficient strains (OD_600_ 0.2) spotted onto LB and LB agar supplemented with 0.2 µg/ml trimethoprim. Images show representative plates from independent triplicate replicates. **c**. Construction of the trimethoprim-resistant *E. coli* background. The promoter region of the *folA* gene (shown as text, with the -35 and -10 regions highlighted in pink) was modified using λ_RED_ recombineering to introduce a C > T SNP (highlighted in orange) in the -35 promoter region. **d**. Spot plate dilution assays of trimethoprim-resistant DNA repair deficient strains, Tmp^R^ (EKW058), Tmp^R^ Δ*recA*::Kan^R^ (EKW059), Tmp^R^ Δ*recB*::Kan^R^ (EKW060), Tmp^R^ Δ*recF*::Kan^R^ (EKW061), Tmp^R^ Δ*recO*::Kan^R^ (EKW062), Tmp^R^ Δ*recR*::Kan^R^ (EKW063) and Tmp^R^ Δ*mutT*::Kan^R^ (EKW064). Dilutions were spotted onto LB and LB agar supplemented with 5 µg/ml trimethoprim. Images show representative plates from independent triplicate replicates.

We then assessed these repair mutants in a trimethoprim-resistant background (Tmp^R^). Resistance to trimethoprim can occur via acquisition of mobile genetic elements or single nucleotide polymorphisms (SNPs). The most common mode of resistance in trimethoprim is SNPs within the drug’s target gene *folA* which encodes for DHFR, or within the *folA* promoter region ^51^. For this study, trimethoprim resistance was conveyed through a clinically relevant SNP (T>C) in the *folA* –35 promoter region (Figure 3c) ^51, 52^. In the Tmp^R^ background, a significant loss of cell viability at 5 µg/ml trimethoprim was observed for cells lacking RecA or RecB (Figure 3d). RecO and RecR deficient mutants showed a slight loss in viability. Thus DSBR mutations increase sensitivity in both antibiotic-sensitive and - resistant backgrounds while SSGR mutants had mixed effects.

### Defects in double-strand break repair reduce tolerance to ampicillin

We next wanted to assess the dependency on DNA repair following treatment with an antibiotic belonging to the β-lactam class. This family of antibiotics target cell wall synthesis. β-lactams block the transpeptidation of peptidoglycan subunits, reducing cell wall integrity, which increase the frequency of cell lysis events ^14^. Here we chose to focus on the β-lactam, ampicillin. Recent studies have demonstrated that treatment of *E. coli* with ampicillin increases cellular ROS levels ^45^, which are proposed to result in the damage of DNA ^47^.

Following MIC analysis, we found that the sensitivity of DNA repair deficient *E. coli* to ampicillin was not significantly altered in comparison to wild type (Supplementary Figure S2a). Minor differences (less than 1 µg/ml) in MIC were observed for Δ*recA* and *ΔrecB* strains, which was complemented *in trans* (Supplementary Figure S2b). This change in MIC is unlikely to be clinically useful. These findings suggest that DNA repair does not play a significant role in bacterial sensitivity following ampicillin treatment.

Ampicillin and other β-lactam antibiotics are more effective during certain bacterial growth phases, particularly stages of high growth ^53^. Delayed growth or dormancy can confer tolerance to ampicillin ^54^. Antibiotic tolerance is a phenotypic phenomenon that transiently increases the resilience of bacterial cells during drug exposure (often at levels much higher than the MIC) prolonging cell survival during treatment ^55^. Importantly, tolerance can also facilitate the evolution of AMR ^56, 57^. To assess effects on tolerance we used a modified version of the TDtest ^58^, measuring the percentage of bacterial regrowth following ampicillin exposure. Cells lacking RecA or RecB demonstrated significantly reduced tolerance to ampicillin, as demonstrated by reduced regrowth of cells within the ZOI following the addition of supplementary nutrients (Figures 4a, 4b and Supplementary Figure 2d). Ampicillin tolerance in *recA* and *recB* mutants was restored to wild-type levels following complementation *in trans* (Figures 4c and 4e). Our findings are in good agreement with previous studies, which have demonstrated that deletion of *recA* reduced the tolerance of *E. coli* to ampicillin during early exposure ^59^.

**Figure 4:**
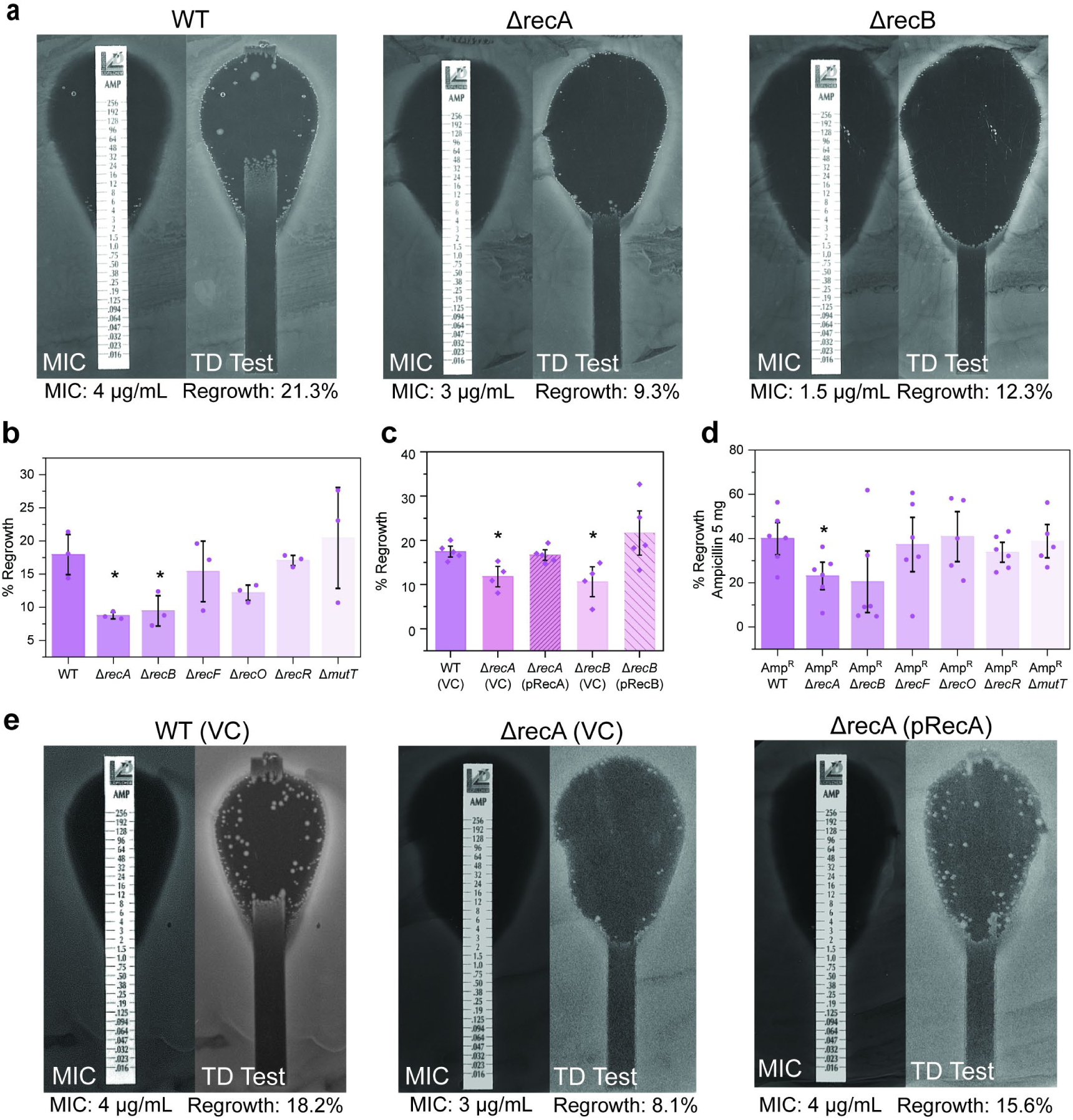
Cells deficient in double-strand break repair have a reduced tolerance to ampicillin. **a.** Representative images of ampicillin MIC and tolerance (TD Test) plate assays for wild-type (WT), Δ*recA* and Δ*recB E. coli* strains. MICs and percentage regrowth are denoted below the images. Images show representative plates from independent triplicate replicates. **b.** Ampicillin percent regrowth values obtained for isogenic *E. coli* strains MG1655 (WT; wild-type), Δ*recA*::Kan^R^ (HH020), Δ*recB*::Kan^R^ (EAW102), Δ*recF*::Kan^R^ (EAW629), Δ*recO*::Kan^R^ (EAW114), Δ*recR*::Kan^R^ (EAW669) and Δ*mutT*::Kan^R^ (EAW999). The means and standard errors of the mean are shown, based on results from at least three biological replicates. Statistical analysis was carried out using a student’s *t*-test. An asterisk denotes statistical significance (*p* < 0.05) compared to wild-type (MG1655). **c.** Ampicillin percent regrowth values obtained for wild-type (MG1655) with empty vector (VC; vector control), Δ*recA* and Δ*recB* mutants with empty vector (VC) and complemented derivatives (pRecA and pRecB, respectively). The means and standard errors of the mean are shown based on results from at least four biological replicates. Statistical analysis was carried out using a student’s *t*-test. An asterisk denotes statistical significance (*p* < 0.05) compared to wild-type empty vector control, WT (VC). **d.** Percent regrowth measurements for ampicillin-resistant WT and DNA repair mutant strains following disk diffusion assays with 5 mg ampicillin. Amp^R^ (COF001), Amp^R^ Δ*recA* (COF002), Amp^R^ Δ*recB* (COF003), Amp^R^ Δ*recF* (COF004), Amp^R^ Δ*recO* (COF005), Amp^R^ Δ*recR* (COF006) and Amp^R^ Δ*mutT* (COF007). The means and standard errors of the mean are shown based on results from at least five biological replicates. Statistical analysis was carried out using a student’s *t*-test. An asterisk denotes statistical significance (*p* < 0.05) compared to the ampicillin-resistant parental strain (Amp^R^). **e.** Representative images of ampicillin MIC and tolerance (TD Test) plate assays for wild-type with empty vector control (WT (VC)), Δ*recA* with empty vector (Δ*recA*(VC)) and complemented (Δ*recA*(pRecA)) derivatives. MICs and percentage regrowth are denoted below the images.

We also observed reduced tolerance for ampicillin-resistant (Amp^R^) Δ*recA* and Δ*recB* cells (Figure 4d and Supplementary Figure 2e). These Amp^R^ strains were constructed by introducing the pWSK29 plasmid (which confers ampicillin resistance through the *bla* gene) into wild-type and DNA repair deficient cells by transformation. Strains deficient in SSGR or NPS had little variation in tolerance to ampicillin in either Amp^S^ or Amp^R^ backgrounds. Our results indicate that the repair of double-strand breaks contributes to ampicillin tolerance in both the ampicillin-sensitive and -resistant backgrounds.

For completeness, we also examined the tolerance phenotypes of the DNA repair mutant strains following exposure to the other drugs used in this study. No tolerant cells were observed following treatment with ciprofloxacin (Supplementary Figure S3a) or kanamycin (Supplementary Figure S3b), this is likely due to the strong bactericidal activity of these antibiotics. Tolerant cells were observed following nitrofurantoin (Supplementary Figure S3c) and trimethoprim (Supplementary Figure S3d) treatment, however no significant changes in the frequency of tolerance were observed for any DNA repair mutant. DNA repair does not appear to play a role in antibiotic tolerance to the drugs nitrofurantoin or trimethoprim.

### DSBR defects supress induction of the SOS response by ciprofloxacin, nitrofurantoin and trimethoprim

The mutagenic SOS response is triggered by some antibiotics ^13^. We qualitatively examined the induction of the SOS response by ciprofloxacin and nitrofurantoin in DNA repair deficient cells using an agar plate-based SOS reporter assay. SOS reporter strains were generated by transformation of DNA repair mutant cells with the plasmid pUA66-P_sulA_-*gfp* ^48^, which places *gfp* under the control of the SOS- inducible promoter P*_sulA_*. When exposed to ciprofloxacin, wild type, Δ*recF,* Δ*recO,* Δ*recR* and Δ*mutT* cells exhibited robust SOS induction, manifesting as a strong fluorescence band at the border of the zone of inhibition (Figure 5a and Supplementary Movie 1). Ciprofloxacin-induced SOS was abolished in the SOS-defective Δ*recA* strain, as expected, as well as the DSBR defective Δ*recB* background. The same pattern of SOS induction, albeit at a reduced intensity and at higher drug concentrations, was also observed in the Cip^R^ background (Figure 5b). This result supports previous findings that SOS induction by quinolones is strongly *recB*-dependent ^32, 60^, and further demonstrated this dependency in a ciprofloxacin-resistant background.

**Figure 5:**
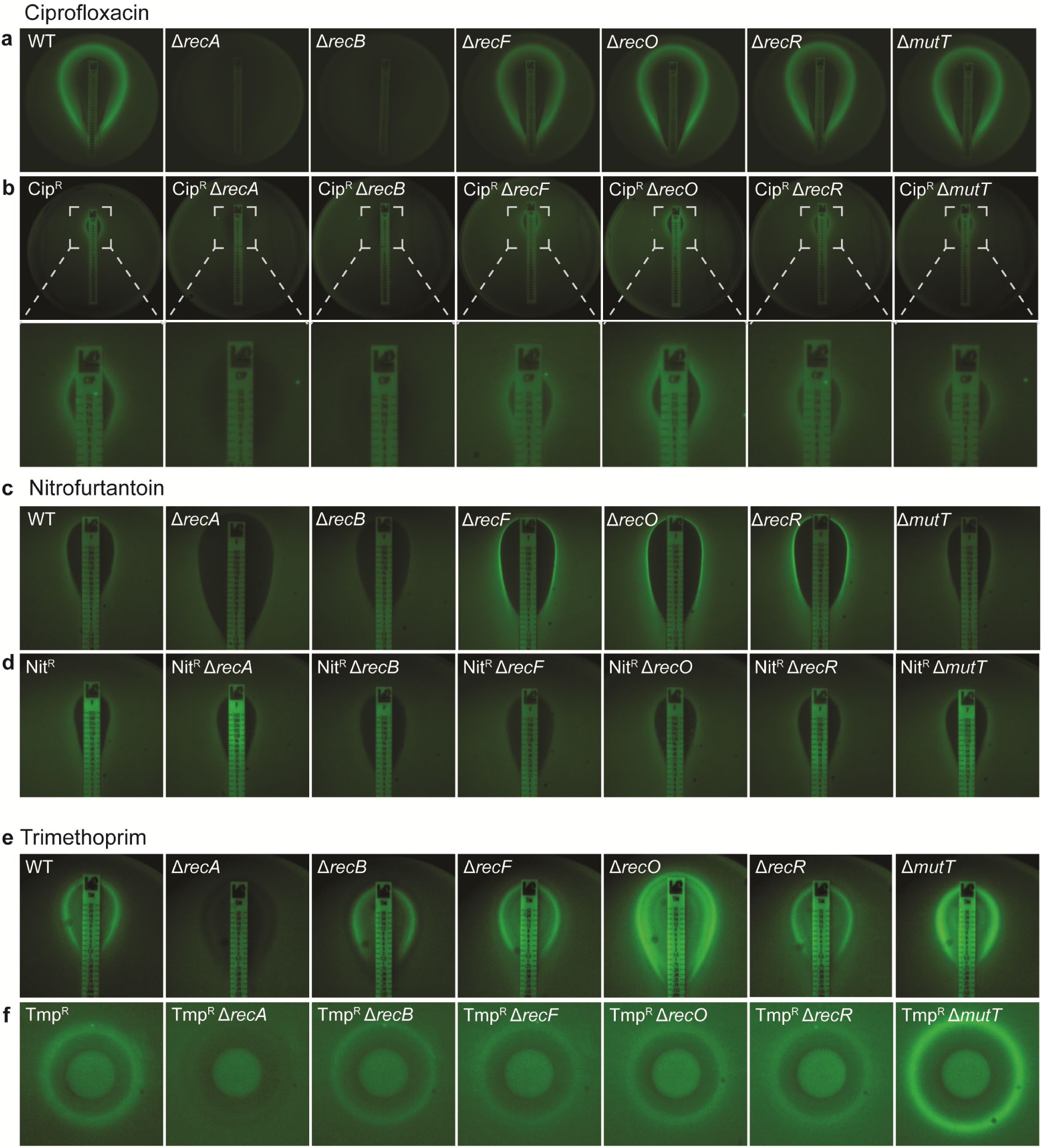
*E.* coli rely on RecB for induction of the SOS response following treatment with ciprofloxacin or nitrofurantoin. **a.** Expression of SOS reporter fusion P*_sulA_*-*gfp* on a solid agar surface in ciprofloxacin-sensitive wild-type (WT) and DNA repair deficient strains grown in the presence of a ciprofloxacin MIC test strip (0.002-32 µg/ml). SOS induction is visualised as strong fluorescence band at the border of the zone of inhibition. **b.** Expression of SOS reporter fusion P*_sulA_*-*gfp* on a solid agar surface in ciprofloxacin-resistant (Cip^R^) wild-type and DNA repair deficient strains grown in the presence of a ciprofloxacin MIC test strip (0.002-32 µg/ml). **c.** Expression of SOS reporter fusion P*_sulA_*-*gfp* on a solid agar surface in antibiotic-sensitive wild-type (WT) and DNA repair deficient strains grown in the presence of a nitrofurantoin MIC test strip (0.032-512 µg/ml). **d.** Expression of SOS reporter fusion P*_sulA_*-*gfp* on a solid agar surface in nitrofurantoin-resistant (Nit^R^) wild-type and DNA repair deficient strains grown in the presence of a nitrofurantoin MIC test strip (0.032-512 µg/ml). **e.** Expression of SOS reporter fusion P*_sulA_*-*gfp* on a solid agar surface in trimethoprim-sensitive wild-type and DNA repair deficient strains grown in the presence of a trimethoprim MIC test strip (0.002-32 µg/ml). **f.** Expression of SOS reporter fusion P*_sulA_*-*gfp* on a solid agar surface in trimethoprim-resistant (Tmp^R^) wild-type (WT) and DNA repair deficient strains grown in the presence of a disk containing 1 mg trimethoprim.

When exposed to nitrofurantoin, wild-type and Δ*mutT* cells containing the SOS-reporter plasmid exhibited weak fluorescence signal (Figure 5c). Strains defective in SOS activation (Δ*recA*) or DSBR (Δ*recB*) resulted in no observable induction of the SOS response fluorescence (Figure 5c). Deletion of SSGR intermediates *recF*, *recO*, and *recR* resulted in high-level SOS induction, as indicated by a bright fluorescence signal at the border of the ZOI and spreading outward (Figure 5c). Thus, deletion of *recA* or DSBR (*recB*) abolishes the SOS response under nitrofurantoin treatment. This would presumably reduce the capacity of these cells to undergo mutagenic repair associated with resistance formation. In contrast, cells lacking SSGR proteins exhibit increased SOS response in response to nitrofurantoin treatment and could potentially become highly mutagenic through this pathway.

When cells become resistant to nitrofurantoin there is a change in the genetics of the SOS response (Figure 5d). In the Nit^R^ background, cells lacking *recO*, *recR* and *mutT* exhibited low-level fluorescence, similar to that of Nit^R^ *rec*^+^ cells. Deletion of *recA* and *recB* abolish the SOS response as consistent with the sensitive background, yet loss of the *recF* gene also abolished SOS in the Nit^R^ background. Thus, in both sensitive and resistant backgrounds, loss of *recA* and DSBR (*recB*) dampens the SOS response under nitrofurantoin treatment. SOS is dependent on *recF* in Nit^S^ cells and independent of *recF* in the Nit^R^ background.

In our hands, no obvious SOS response was detected in any strain following ampicillin treatment up to concentrations of 256 μg/mL (Supplementary Figure S2d). This finding contrasts with previous studies ^17, 61^ which have demonstrated ampicillin-dependent SOS induction in *E. coli* using similar methods. We observed that kanamycin treatment did not induce a detectable SOS response in any strain analysed (Supplementary Figure S1d). In agreement with other studies ^10, 17, 59^ kanamycin treatment does not elicit a detectable SOS response in *E. coli*. Trimethoprim did induce a clear SOS response signal in both the sensitive and resistant backgrounds (Figures 5e and 5d). Disruption of *recA* eliminated SOS response in both cases. Deletion of *recB* reduced SOS in both cases. Disruption of SSGR did not reduce SOS, except for a *recO* deletion in the Tmp^R^ background. We note that SOS signal was enhanced in the *recO* deletion in the sensitive background.

### Putative DSBR inhibitors ML328 and IMP-1700 exhibit off-target effects

We next examined two putative DSBR inhibitors that have been reported in the literature, ML328 ^62^ and IMP-1700 ^63^. In these studies, the two compounds demonstrated specific affinities to the *E. coli* RecBCD complex or the functionally related AddAB(RexAB) complexes from *Helicobacter pylori* and *Staphylococcus aureus.* Additionally, IMP-1700 was claimed to potentiate ciprofloxacin activity, sensitising a multi-drug resistant *S. aureus* strain to clinically relevant levels of ciprofloxacin ^63^. Whilst these two compounds show early promise, further work is required to confirm the mechanism of action as being inhibition of DSBR.

We examined the biological activity of these two drugs in DNA-repair deficient *E. coli* derivatives using disc diffusion assays (Figures 6a and 6d). We expected that cells lacking DSBR should not show sensitivity to these compounds, since the drug target (RecB) was no longer present. However, we observed that deletion of *recB* resulted in an increased sensitivity to both drugs, suggesting that there may be off target effects. Cells lacking *recA* were also significantly more sensitive to both compounds. Complementation of *recA* and *recB* mutants returned sensitivity to both drugs back to wild-type levels, confirming the roles of RecA and RecB in survival following ML328 and IMP-1700 treatment (Figures 6b and 6e). We quantitatively determined the MICs of ML328 and IMP-1700 in wild-type, Δ*recA* and Δ*recB* mutants. Consistent with the disc diffusion data, deletion of *recA* or *recB* reduced the MICs of both compounds in *E. coli* (Table 1 and Supplementary Figure S5).

**Figure 6:**
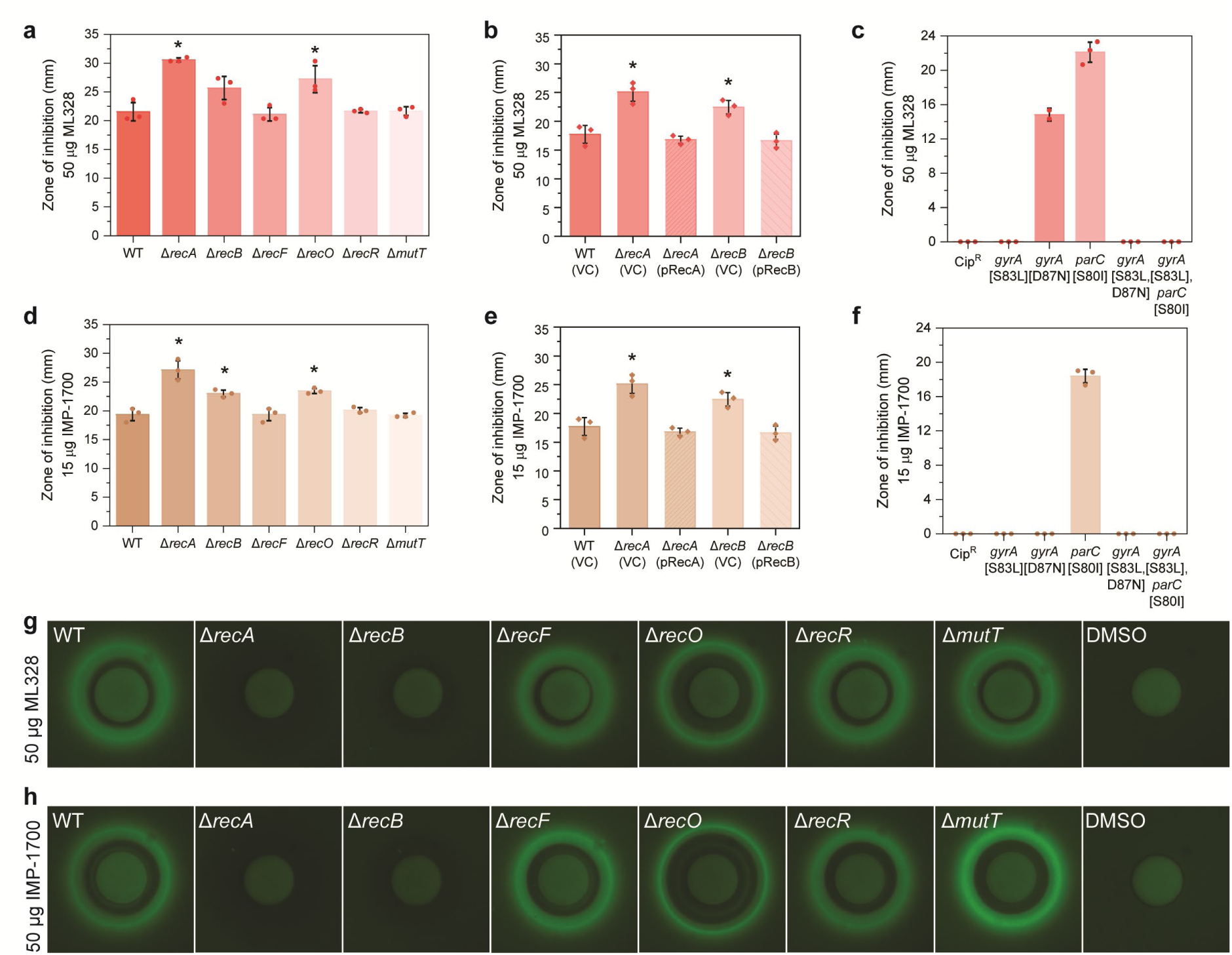
Cells lacking RecA and RecB are more sensitive than wild-type to ML328 and IMP-1700. **(a. and d.)** Zone of inhibition area measurements for isogenic *E. coli* strains MG1655 (WT; wild-type), Δ*recA*::Kan^R^ (HH020), Δ*recB*::Kan^R^ (EAW102), Δ*recF*::Kan^R^ (EAW629), Δ*recO*::Kan^R^ (EAW114), Δ*recR*::Kan^R^ (EAW669) and Δ*mutT*::Kan^R^ (EAW999) following disk diffusion assays with (**a**) 50 µg ML328 or (**d**) 15 µg IMP-1700. The means and standard errors of the mean are shown based on results from at least three biological replicates. Statistical analysis was carried out using a student’s *t*-test. An asterisk denotes statistical significance (*p* < 0.05) compared to wild-type (WT). **(b. and e.)** Zone of inhibition area values obtained for wild-type (MG1655) with empty vector (VC; vector control), Δ*recA* and Δ*recB* mutants with empty vector and complemented derivatives (pRecA and pRecB, respectively) following disk diffusion assays with (**b**) 50 µg ML328 or (**e**) 15 µg IMP-1700. The means and standard errors of the mean are shown based on results from at least three biological replicates. Statistical analysis was carried out using a student’s *t*-test. An asterisk denotes statistical significance (*p* < 0.05) compared to wild-type empty vector control (WT (VC)). **(c. and f.)** Zone of inhibition area values obtained for isogenic *E. coli* derivatives possessing single, double or triple point mutations that confer ciprofloxacin resistance following disk diffusion assays with (**c**) 50 µg ML328 or (**f**) 15 µg IMP-1700. Strains used are Cip^R^ (CH5741; *gyrA;* [S83L, D87N] and *parC;* [S80I]), *gyrA* [S83L] (LM328), *gyrA;* [D87N] (LM534), *parC;* [S80I] (LM792), *gyrA;* [S83L, D87N] (LM625) and *gyrA;* [S83L] *parC;* [S80I] (CH6179). The means and standard errors of the mean are shown based on results from at least three biological replicates. **(g. and h.)** Expression of SOS reporter fusion P*_sulA_*-*gfp* on a solid agar surface in wild-type (WT) and DNA repair deficient strains grown in the presence of (**g**) 50 µg ML328 or (**h**) 50 µg IMP-1700. SOS induction is visualised as strong fluorescence band at the border of the zone of inhibition.

**Table 1:** MIC and IC_50_ values for ML328 and IMP-1700.

| Strain | ML328 |  | IMP-1700 |  |
| --- | --- | --- | --- | --- |
|  | MIC (μg/ml) | IC<sub>50</sub> (μg/ml) | MIC (μg/ml) | IC<sub>50</sub> (μg/ml) |
| WT | 0.5 ± 0.00 | 0.255 ± 0.03 | 1 ± 0.00 | 0.481 ± 0.05 |
| *ΔrecA* | 0.125 ± 0.00 | 0.036 ± 0.002 | 0.125 ± 0.00 | 0.032 ± 0.003 |
| *ΔrecB* | 0.167 ± 0.07 | 0.036 ± 0.003 | 0.25 ± 0.00 | 0.049 ± 0.003 |

IC<sub>50</sub> values were determined by dose-response non-linear regression. Data represent the mean and standard error based on three biological replicates.

Lim, et al. ^63^ showed inhibition of the SOS response following treatment with IMP-1700. In contrast, we observed direct induction of the SOS response by these compounds in *E. coli* (Figures 6g and 6h). For both compounds, SOS induction was abolished in the SOS-defective Δ*recA* strain as expected, and absent in the DSBR-defective Δ*recB* background. No significant changes in SOS induction were observed in wild-type, *recF, recO, recR* or *mutT* mutants. These results suggest that induction of the SOS response in *E. coli* following treatment with ML328 and IMP-1700 is dependent on RecB.

Both the DSBR-inhibiting compounds ML328 and IMP-1700 were constructed using quinolone structural backbones. We reasoned that they each might inhibit DNA gyrase and topoisomerase IV. The authors ^63^ examined this potential activity using purified proteins in bulk biochemical assays and claimed no significant interactions. However, the patterns of strain sensitivity we observed with these two compounds and the strong *recB*-dependent SOS induction mimicked the results we had obtained for ciprofloxacin. We reasoned that if ML328 and IMP-1700 inhibited DNA gyrase and topoisomerase IV, mutations conferring resistance to quinolones may also confer resistance to these two compounds. We repeated the disc diffusion assays using Cip^R^ DNA repair deficient derivatives (Supplementary Figures S6a and S6b). All strains examined were found to be resistant to both ML328 and IMP-1700 at the concentrations tested. Using broth micro dilution assays, the MICs of the Cip^R^ strain for both compounds were determined to be greater than 128 µg /ml (Supplementary Figures S6c and S6d). These findings show that the point mutations *gyrA;* [S83L, D87N] and *parC;* [S80I], which typically confer resistance to quinolones, also confer resistance to ML328 and IMP-1700. To confirm which of the three point-mutations was most important for conferring resistance to these compounds, we repeated disc diffusion assays with isogenic *E. coli* derivatives possessing one single point mutation, or two mutations in combination (Figures 6c and 6f). Cells possessing a single S80I point mutation in *parC* were sensitive to both ML328 and IMP-1700, suggesting topoisomerase is not the primary target for these two drugs. The single point mutation D87N in *gyrA* conferred resistance to IMP-1700, but not ML328, however the single point mutation S83L in *gyrA* conferred resistance to both compounds. These findings suggest that whilst these two compounds may target DSBR to some degree, in our hands the primary mode of action in *E. coli* is inhibition of gyrase and topoisomerase IV.

## Discussion

This study demonstrated that disruption of bacterial DSBR induced sensitising effects against multiple bactericidal antibiotics with disparate modes of action. The deletion of genes involved in DSBR (*recA* and *recB*) hypersensitised *E. coli* cells against ciprofloxacin and nitrofurantoin (Figure 1), enhanced the clearing of cells by kanamycin (Figure 2) and trimethoprim (Figure 3), and decreased tolerance to ampicillin (Figure 4). Most importantly, sensitising effects were also observed in antibiotic-resistant strains, raising the possibility of targeting DSBR for the development of novel resistance breakers. Additionally, activation of the highly mutagenic SOS response was found to be dependent on DSBR, raising the possibility that disrupting DSBR would limit the capacity of bacterial populations to develop further antibiotic resistance mutations. Overall, our findings establish DSBR as a promising target for the design of broad-range resistance-breaking compounds that could be used to dramatically enhance the effectiveness of existing bactericidal antibiotics and to supress the development of antibiotic resistance.

### Double-strand break repair as a novel drug target

In this study, we found that disruption of DSBR induced a suite of phenotypes that could lead to more effective antibiotic treatments. Disruption of DSBR enhanced killing by five disparate classes of antibiotics. Previous studies have shown that *recA* mutants are highly sensitive to quinolones and other DNA damaging drugs ^17, 64–66^. Our observations support and build upon these findings, demonstrating that *recB* mutants also share this hyper-sensitivity phenotype. Importantly, we observed that these sensitisation phenomena also extended to drug-resistant *E. coli* strains, suggesting for the first time that disruption of DSBR via inhibition of RecA or RecBCD might represent a viable strategy for the development of broad-ranging antibiotic resistance breakers.

In this work, we examined *E. coli*, however studies by others suggests that disruption of DSBR may improve the killing of other bacterial species. In *Staphylococcus aureus*, DSBR pathways promote the survival of both antibiotic-sensitive and -resistant bacteria following exposure to antibiotics such as fluoroquinolones, daptomycin and nitrofurantoin ^67^. In a separate study, DSBR-deficient *Acinetobacter baumannii* were significantly sensitised to colistin, gentamycin, rifampicin and tigecycline ^68^. For both pathogens, inactivation of DSBR pathways also increased susceptibility to trimethoprim and sulfamethoxazole ^69, 70^. Importantly, DSBR pathways also play significant roles in the establishment and maintenance of *S. aureus* infections ^71^. This reliance on DSBR for infection is also true for the non-ESKAPE pathogens *Helicobacter pylori, Salmonella enterica* and *Campylobacter jejuni* ^72–74^. There are few exceptions to this finding, for example, DSBR-deficient strains of the acid-fast bacterium *Mycobacterium tuberculosis* do not have any infectivity defects ^75^. However, this is likely due to the presence of alternate pathways that can repair double-strand DNA breaks in these cells^76^. Regardless, these results in conjunction with our own findings, suggest that DSBR inhibitors may not only potentiate antibiotic activities, but also reduce the infection potential of many diverse bacterial pathogens.

At high concentrations, the aminoglycoside kanamycin and the folate inhibitor trimethoprim were more active against *recA* and *recB* mutants than the wild-type strain. These sensitivities were also observed in the respective resistant backgrounds. It remains unclear why the MICs of DSBR-deficient strains were similar to wild-type whilst also demonstrating notably increased sensitivities to high drug concentrations. One possibility is that high concentrations of these drugs are necessary for initiating DNA damage. Although disruption of DSBR does not significantly re-sensitise *E. coli* to these drugs, it does increase the efficacy of killing. It is reasonable to hypothesise that the increased killing observed here may translate to improved infection clearance times *in vivo* when using combinational DSBR inhibitor-antibiotic therapeutics.

In both the sensitive and resistant backgrounds we observed a strong dependence on both RecA and RecB for tolerance following ampicillin exposure. Tolerance enables bacteria to survive exposure to high levels of antibiotic ^55^. Importantly, antibiotic tolerance is a key contributor to recalcitrant infections, since these cells survive primary antibiotic treatment ^77^. Antibiotic tolerance can also act as a precursor for resistance development ^57^. Our discovery that ampicillin tolerance in *E. coli* relies on DSBR suggests that combinational therapy with DSBR-inhibiting drugs may help to eliminate tolerant bacteria.

Mutagenesis is one of the major pathways through which antibiotic resistance develops in bacteria ^13, 28^. In DNA-damage settings, including certain antibiotic treatments, the rate of mutagenesis is elevated through activation of the SOS response ^28^. It is at concentrations of drug higher than the MIC (where often the SOS response is activated) but within a concentration range where cells are not yet effectively killed, that the development and subsequent selection for drug-resistant mutants frequently occurs. This antibiotic concentration range is known as the mutant selection window ^12^. For treatments to be effective at preventing the development of mutational resistance, antibiotic concentrations need to be maintained above this mutant selection window. Alternatively, strategies could prevent mutagenesis altogether, through suppression of the SOS response. Disrupting DSBR does both. Recent findings have linked ciprofloxacin-induced SOS activation with DSBR ^32^. Our observations are in good agreement with this study. Further, our work has established a new link between DSBR and nitrofurantoin-induced SOS induction, since *recB* mutants did not elicit an SOS response to nitrofurantoin. Importantly, this RecB dependence for SOS induction also applies in drug-resistant *E. coli* for both ciprofloxacin and nitrofurantoin. We hypothesise that cells unable to initiate SOS, or have a reduced induction of SOS, would be less likely to develop beneficial resistance mutations. Inhibition of SOS induction through disruption of DNA repair, specifically DSBR, may prove a promising strategy to combat the evolution of antimicrobial resistance.

### Multiple bactericidal antibiotics induce double-strand breaks

It is well characterised that ciprofloxacin-induced DNA damage predominantly occurs in the form of double-stranded breaks ^20, 32, 33^. For the other antibiotics used in this study, the mechanisms that drive DNA damage remain largely unknown. Throughout this study, we observed several antibiotic-associated phenotypes that were dependent on DSBR. Kanamycin and trimethoprim treatment at high concentrations was more effective at clearing cells lacking DSBR than wild-type, suggesting at high drug concentrations double-strand DNA breaks occur. We also observed reduced ampicillin tolerance in DSBR deficient strains, suggesting that long-term exposure to ampicillin induces double-stranded breaks. In the case of nitrofurantoin, survival of cells following antibiotic treatment was strongly dependent on homologous recombination, including the DSBR pathway. This finding suggests that, in part, that the formation of double-stranded breaks is one mechanism by which nitrofurantoin works in *E. coli*. Taken together, this study lends further weight to the notion that DNA damage, and in particular double-strand breaks, is a common thread between many bactericidal antibiotics.

### New antibiotic induced phenotypes associated with DNA gap repair

Single-strand DNA gaps in bacteria are commonly formed post-replication ^15^, or via exposure to DNA damaging agents, such as UV ^78^ and possibly even antibiotics. If left unrepaired, these gaps can be converted into DSBs, which are highly detrimental to bacterial cells ^50, 79^. Although it was not observed for all antibiotics tested in this study, the deletion of components involved in SSGR (*recF, recO* and *recR*) did significantly alter some antibiotic sensitivity phenotypes. Notably, all mutants lacking components of SSGR were sensitised to nitrofurantoin, to levels equal to the DSBR-deficient cells. This sensitisation effect in SSGR-deficient cells was also observed in the nitrofurantoin-resistant background. This suggests that DNA damage in the form of both single-stranded gaps and double-strand breaks are key contributors to nitrofurantoin action in *E. coli*. One unanticipated finding related to the genetic dependencies of SOS induction in cells exposed to nitrofurantoin. MICs were significantly reduced in SSGR-deficient strains, yet these cells demonstrated high levels of SOS relative to the wild type. This increased-SOS induction behaviour was not observed in Nit^R^ cells. In the case of nitrofurantoin, targeting single-cell gap repair does increase antibiotic sensitivity, however it may also enhance the likelihood of resistant cells forming due elevated SOS activity.

For the drugs ciprofloxacin and trimethoprim, antibiotic-sensitive *recO* cells were significantly sensitised whilst other SSGR mutants remained largely unchanged. Interestingly, the RecO protein is known to bind both single-stranded DNA and double-stranded DNA ^80^. We also saw that these RecO deficient cells exhibited an increased relative SOS signal during exposure to trimethoprim. However, for both drugs, this sensitisation effect (and for trimethoprim the SOS effect) was no longer present in their respective drug-resistant backgrounds. While further work is required to understand the basis of these phenotypes, they do lend further support to the notion that the role of RecO in SSGR may be somewhat separate from those played by RecF and RecR ^81^.

We also examined the importance of NPS in survival following antibiotic treatment. Antibiotic-induced oxidative stress can result in the formation of highly toxic and mutagenic oxidised nucleotide bases (eg. 8-oxo-dGTP) ^46^, which must be cleared from the nucleotide pool by MutT before they are incorporated into the genome ^27^. Although MTH1 (the MutT homologue in humans) holds promise as an anti-cancer therapeutic target ^82^, disruption of *mutT* does not appear to be sufficiently important for bacterial cell survival or tolerance to be clinically useful. Furthermore, bacterial cells lacking *mutT* are highly mutagenic ^27^, and following trimethoprim treatment, *mutT* mutants had an increased relative induction of the mutagenic SOS response. As such, MutT would not be an appropriate target for future drug development.

### Development of DSBR inhibitors: the challenge of off-target effects

Since our work demonstrated multiple benefits of targeting DSBR we investigated the efficacy of the published DSBR inhibitors ML328 ^62^ and IMP-1700 ^63^ in *E. coli*. Contrary to expectations, cells lacking RecB (the putative target of these compounds) were more sensitive to these two compounds than wild-type. This result implied that there these compounds have off-target effects in *E. coli.* Further investigation revealed that these drugs were rendered ineffective in fluoroquinolone-resistant derivatives of *E. coli*. The putative primary target of these two drugs appears to be DNA gyrase, as a single point mutation (known to increase ciprofloxacin resistance) in *gyrA* rendered cells resistant to these compounds. In the initial studies on these compounds, bulk biochemical assays determined there was no notable inhibition of *E. coli* DNA gyrase or topoisomerase IV activity ^63^. However, only a single concentration of inhibitor was used to assess this inhibition activity. It is likely if the drug concentration range were extended further, gyrase inhibition may have been observed.

Clearly, further work is needed to identify DSBR inhibitors with fewer off-target effects. Nevertheless, the results of the current study highlight that disrupting bacterial DSBR produces multiple beneficial effects and the search for DSBR inhibitors is a worthwhile pursuit.

## Materials and methods

### Bacterial strains, plasmids and culture conditions

*Escherichia coli* strains and plasmids used in this study are listed in Tables 2 and 3 respectively. *E. coli* was cultured at 37°C in Luria-Bertani (BD) broth or agar (1.5% (w/v); BD). As required, media were supplemented with antibiotics (Sigma) at the following concentrations unless otherwise stated kanamycin (Kan; 25 µg/ml), ampicillin (Amp; 100 µg/ml), trimethoprim (Trp; 50 µg/ml), spectinomycin (Spec; 50 µg/ml).

**Table 2:** E. coli strains used in this study

| **Strain** | **Relevant Genotype** | **Parent Strain** | **Source/ Technique** |
| --- | --- | --- | --- |
| MG1655 | *recA<sup>+</sup> recB<sup>+</sup> recF<sup>+</sup> recO<sup>+</sup> recR<sup>+</sup> mutT<sup>+</sup>* | - |  |
| HH020 | *ΔrecA::Kan<sup>R</sup>* | **MG1655** | 85 |
| EAW102 | *ΔrecB::Kan<sup>R</sup>* | **MG1655** | 32 |
| EAW629 | *ΔrecF::Kan<sup>R</sup>* | **MG1655** | 86 |
| EAW114 | *ΔrecO::Kan<sup>R</sup>* | **MG1655** | 86 |
| EAW669 | *ΔrecR::Kan<sup>R</sup>* | **MG1655** | 86 |
| EAW999 | *ΔmutT::Kan<sup>R</sup>* | **MG1655** | Lambda Red recombination |
| HH021 | *ΔrecA::FRT* | **HH020** | 91 |
| HG356 | *ΔrecB::FRT* | **EAW102** | 32 |
| MV009 | *ΔrecF::FRT* | **EAW629** | This work |
| SRM019 | *ΔrecO::FRT* | **EAW114** | This work |
| SRM020 | *ΔrecR::FRT* | **EAW669** | This work |
| MV005 | *ΔmutT::FRT* | **EAW999** | This work |
| CH5741 | Cip<sup>R</sup> (*gyrA* [S83L, D87N] *parC* [S80I]) *recA*<sup>+</sup> *recB*<sup>+</sup> *recF*<sup>+</sup> *recO*<sup>+</sup> *recR*<sup>+</sup> *mutT* | **MG1655** | 38 |
| FM002 | Cip<sup>R</sup> (*gyrA* [S83L, D87N] *parC* [S80I]) $\Delta recA::Kan^R$ | **CH5741** | Transduction of CH5741 with P1 grown on HH021 |
| FM001 | Cip<sup>R</sup> (*gyrA* [S83L, D87N] *parC* [S80I]) $\Delta recB::Kan^R$ | **CH5741** | Transduction of CH5741 with P1 grown on EAW102 |
| FM003 | Cip<sup>R</sup> (*gyrA* [S83L, D87N] *parC* [S80I]) $\Delta recF::Kan^R$ | **CH5741** | Transduction of CH5741 with P1 grown on EAW629 |
| FM004 | Cip<sup>R</sup> (*gyrA* [S83L, D87N] *parC* [S80I]) $\Delta recO::Kan^R$ | **CH5741** | Transduction of CH5741 with P1 grown on EAW114 |
| FM005 | Cip<sup>R</sup> (*gyrA* [S83L, D87N] *parC* [S80I]) $\Delta recR::Kan^R$ | **CH5741** | Transduction of CH5741 with P1 grown on EAW669 |
| MV001 | Cip<sup>R</sup> (*gyrA* [S83L, D87N] *parC* [S80I]) $\Delta mutT::Kan^R$ | **CH5741** | Transduction of CH5741 with P1 grown on EAW999 |
| SSH021 | MG1655 (pJM1071) | **MG1655** | Transformation of MG1655 with pJM1071 |
| SRM009 | $\Delta recA::Kan^R$ (pJM1071) | **HH020** | Transformation of HH020 with pJM1071 |
| MV002 | $\Delta recA::Kan^R$ (pRecA) | **HH020** | Transformation of EAW102 with pRecA |
| SRM010 | $\Delta recB::Kan^R$ (pJM1071) | **EAW102** | Transformation of EAW102 with pJM1071 |
| EKW008 | $\Delta recB::Kan^R$ (pRecB) | **EAW102** | Transformation of EAW102 with pRecB |
| SRM011 | $\Delta recA::FRT$ (pJM1071) | **HH021** | Transformation of HH021 with pJM1071 |
| CD006 | $\Delta recA::FRT$ (pRecA) | **HH021** | Transformation of HH021 with pRecA |
| SRM012 | $\Delta recB::FRT$ (pJM1071) | **HG356** | Transformation of HG356 with pJM1071 |
| SRM013 | $\Delta recB::FRT$ (pRecB) | **HG356** | Transformation of HG356 with pRecB |
| COF001 | Amp<sup>R</sup> MG1655 (pWSK29) | **MG1655** | Transformation of MG1655 with pWSK29 |
| COF002 | Amp<sup>R</sup> $\Delta recA::Kan^R$ (pWSK29) | **HH020** | Transformation of HH020 with pWSK29 |
| COF003 | Amp<sup>R</sup> $\Delta recB::Kan^R$ (pWSK29) | **EAW102** | Transformation of EAW102 with pWSK29 |
| COF004 | Amp<sup>R</sup> $\Delta recF::Kan^R$ (pWSK29) | **EAW629** | Transformation of EAW629 with pWSK29 |
| COF005 | Amp<sup>R</sup> $\Delta recO::Kan^R$ (pWSK29) | **EAW114** | Transformation of EAW114 with pWSK29 |
| COF006 | Amp<sup>R</sup> $\Delta recR::Kan^R$ (pWSK29) | **EAW669** | Transformation of EAW669 with pWSK29 |
| COF007 | Amp<sup>R</sup> $\Delta mutT::Kan^R$ (pWSK29) | **EAW999** | Transformation of EAW999 with pWSK29 |
| CD001 | MG1655 (pUA66) | **MG1655** | Transformation of MG1655 with pUA66 |
| CD002 | Kan<sup>R</sup> $\Delta recA::FRT$ (pUA66) | **HH021** | Transformation of HH021 with pUA66 |
| CD003 | Kan<sup>R</sup> $\Delta recB::FRT$ (pUA66) | **HG356** | Transformation of HG356 with pUA66 |
| CD004 | Kan<sup>R</sup> $\Delta recF::FRT$ (pUA66) | **MV009** | Transformation of MV009 with pUA66 |
| SRM026 | Kan<sup>R</sup> $\Delta recO::FRT$ (pUA66) | **SRM019** | Transformation of SRM019 with pUA66 |
| SRM027 | Kan<sup>R</sup> $\Delta recR::FRT$ (pUA66) | **SRM020** | Transformation of SRM020 with pUA66 |
| CD005 | Kan<sup>R</sup> $\Delta mutT::FRT$ (pUA66) | **MV005** | Transformation of MV005 with pUA66 |
| SSH091 | MG1655 (pUA66-$P_{sulA}$-*gfp*) | **MG1655** | <sup>32</sup> |
| MV003 | $\Delta recA::FRT$ (pUA66-$P_{sulA}$-*gfp*) | **HH021** | Transformation of HH021 with pUA66-$P_{sulA}$-*gfp* |
| SSH111 | $\Delta recB::FRT$ (pUA66-$P_{sulA}$-*gfp*) | **HG356** | <sup>32</sup> |
| MV011 | $\Delta recF::FRT$ (pUA66-$P_{sulA}$-*gfp*) | **MV009** | Transformation of MV009 with pUA66-$P_{sulA}$-*gfp* |
| SRM028 | $\Delta recO::FRT$ (pUA66-$P_{sulA}$-*gfp*) | **SRM019** | Transformation of SRM019 with pUA66-$P_{sulA}$-*gfp* |
| SRM029 | $\Delta recR::FRT$ (pUA66-$P_{sulA}$-*gfp*) | **SRM020** | Transformation of SRM020 with pUA66-$P_{sulA}$-*gfp* |
| MV013 | $\Delta mutT::FRT$ (pUA66-$P_{sulA}$-*gfp*) | **MV005** | Transformation of MV005 with pUA66- $P_{sulA}$-*gfp* |
| SRM020 | MG1655 (pEAW915) | **MG1655** | Transformation of MG1655 with pEAW915 <sup>92</sup> |
| SRM037 | $\Delta recA::FRT$ (pEAW915) | **HH021** | Transformation of HH021 with pEAW915 |
| SRM021 | $\Delta recB::FRT$ (pEAW915) | **HG356** | Transformation of HG356 with pEAW915 |
| SRM022 | $\Delta recF::FRT$ (pEAW915) | **MV009** | Transformation of MV009 with pEAW915 |
| SRM023 | $\Delta recO::FRT$ (pEAW915) | **SRM019** | Transformation of SRM019 with pEAW915 |
| SRM024 | $\Delta recR::FRT$ (pEAW915) | **SRM020** | Transformation of SRM020 with pEAW915 |
| SRM025 | $\Delta mutT::FRT$ (pEAW915) | **MV005** | Transformation of MV005 with pEAW915 |
| JW0835-1 | $\Delta nfsA::kan^R$ | **BW25113** | <sup>88</sup> |
| EKW036 | Nit<sup>R</sup> MG1655 $\Delta nfsA$ | **MG1655** | Transduction of MG1655 with P1 grown on JW0835-1 |
| EKW037 | Nit<sup>R</sup> $\Delta recA::FRT \Delta nfsA$ | **HH021** | Transduction of HH021 with P1 grown on JW0835-1 |
| EKW038 | Nit<sup>R</sup> $\Delta recB::FRT \Delta nfsA$ | **HG356** | Transduction of HG356 with P1 grown on JW0835-1 |
| EKW039 | Nit<sup>R</sup> $\Delta recF::FRT \Delta nfsA$ | **MV009** | Transduction of MV009 with P1 grown on JW0835-1 |
| EKW040 | Nit<sup>R</sup> $\Delta recO::FRT \Delta nfsA$ | **SRM019** | Transduction of SRM019 with P1 grown on JW0835-1 |
| EKW041 | Nit<sup>R</sup> $\Delta recR::FRT \Delta nfsA$ | **SRM020** | Transduction of SRM020 with P1 grown on JW0835-1 |
| EKW042 | Nit<sup>R</sup> $\Delta mutT::FRT \Delta nfsA$ | **MV005** | Transduction of MV005 with P1 grown on JW0835-1 |
| EKW058 | Tmp<sup>R</sup> MG1655 [C49765T] | **MG1655** | $\lambda_{RED}$ recombination of MG1655 with SRP84 |
| EKW059 | Tmp<sup>R</sup> $\Delta recA::Kan^R$ | **EKW048** | Transduction of EKW048 with P1 grown on HH020 |
| EKW060 | Tmp<sup>R</sup> $\Delta recB::Kan^R$ | **EKW048** | Transduction of EKW048 with P1 grown on EAW102 |
| EKW061 | Tmp<sup>R</sup> $\Delta recF::Kan^R$ | **EKW048** | Transduction of EKW048 with P1 grown on EAW629 |
| EKW062 | Tmp<sup>R</sup> $\Delta recO::Kan^R$ | **EKW048** | Transduction of EKW048 with P1 grown on EAW114 |
| EKW063 | Tmp<sup>R</sup> $\Delta recR::Kan^R$ | **EKW048** | Transduction of EKW048 with P1 grown on EAW669 |
| EKW064 | Tmp<sup>R</sup> $\Delta mutT::Kan^R$ | **EKW048** | Transduction of EKW048 with P1 grown on EAW999 |
| LM378 | *gyrA* [S83L], *recA*<sup>+</sup> *recB*<sup>+</sup> *recF*<sup>+</sup> *recO*<sup>+</sup> *recR*<sup>+</sup> *mutT*<sup>+</sup> | **MG1655** | 93 |
| LM534 | *gyrA* [D87N], *recA*<sup>+</sup> *recB*<sup>+</sup> *recF*<sup>+</sup> *recO*<sup>+</sup> *recR*<sup>+</sup> *mutT*<sup>+</sup> | **MG1655** | 93 |
| LM625 | *gyrA* [S83L, D87N], *recA*<sup>+</sup> *recB*<sup>+</sup> *recF*<sup>+</sup> *recO*<sup>+</sup> *recR*<sup>+</sup> *mutT*<sup>+</sup> | **MG1655** | 93 |
| LM792 | *parC* [S80I], *recA*<sup>+</sup> *recB*<sup>+</sup> *recF*<sup>+</sup> *recO*<sup>+</sup> *recR*<sup>+</sup> *mutT*<sup>+</sup> | **MG1655** | 93 |
| CH6179 | *gyrA* [S83L], *parC* [S80I], *recA*<sup>+</sup> *recB*<sup>+</sup> *recF*<sup>+</sup> *recO*<sup>+</sup> *recR*<sup>+</sup> *mutT*<sup>+</sup> | **MG1655** | 38 |

**Table 3:** Plasmids used in this study

| Plasmid | Description | Source |
| --- | --- | --- |
| pUA66 | *E. coli* low copy number vector, pSC101 *ori*, GFP reporter plasmid carrying *gfpmut2*, no promoter Kan<sup>R</sup> | 48 |
| pUA66-P<sub>*sulA*</sub>-*gfp* | GFP reporter plasmid carrying *gfpmut2*, *sulA* promoter, Kan<sup>R</sup> | 48 |
| pEAW915 | pACYC184 base, p15A *ori*, GFP reporter plasmid carrying supergloGFP (from pQBI63) behind *recN* (+200 to -21) promoter, Cm<sup>R</sup> | 92 |
| pWSK29 | *E. coli* low copy number vector, pSC101 *ori*, Amp<sup>R</sup>, | 94 |
| pJM1071 | *E. coli* base plasmid, pSC101 *ori*, Spec<sup>R</sup> |  |
| pHG134 (pRecA) | RecA complementation plasmid, *recA* cloned in pJM1071 between NdeI/XbaI, Spec<sup>R</sup> | 85 |
| pSRM3 (pRecB) | RecB complementation plasmid, *recB* plus 200bp upstream cloned in pJM1071 between KpnI/XbaI, Spec<sup>R</sup> | This study |
| pLH29 | FLP expression plasmid, p15A *ori*, FLP recombinase under control of *lacZ* promoter, Amp<sup>R</sup> | 87 |
| pKD46 | Temperature-sensitive $\lambda$ Red recombinase expression plasmid, pSC101 *ori*, rep101ts, Amp<sup>R</sup> | 84 |

### Molecular techniques

Plasmid DNA was extracted from *E. coli* using QIAprep Spin Miniprep kits (Qiagen) as outlined by the manufacturer. *E. coli* cells were made competent and transformed as previously described ^83^. Oligonucleotides used in this study are listed in Table 4 and were synthesised by Integrated DNA Technologies (IDT). PCR amplification was performed using Quick*Taq* (Roche) as recommended by the manufacturer.

**Table 4:**
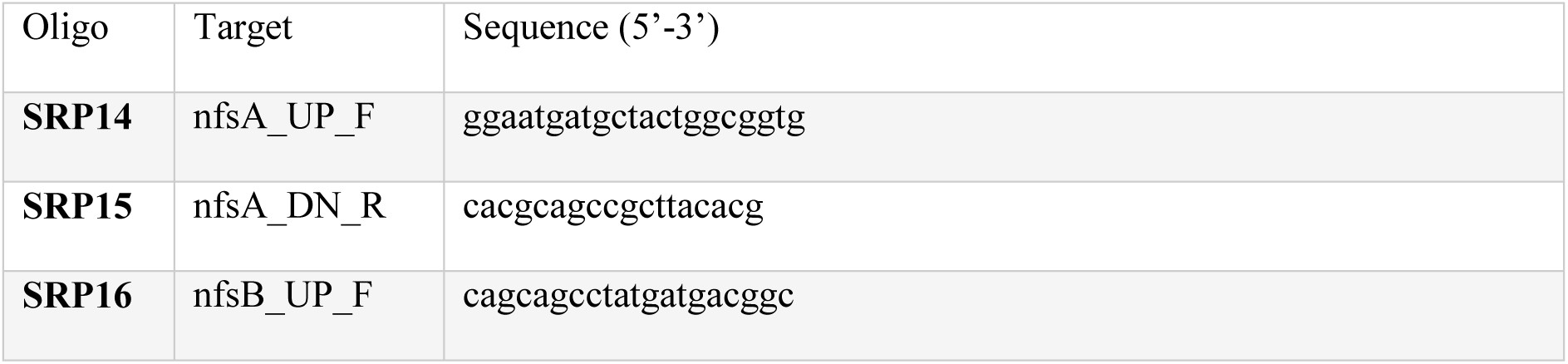
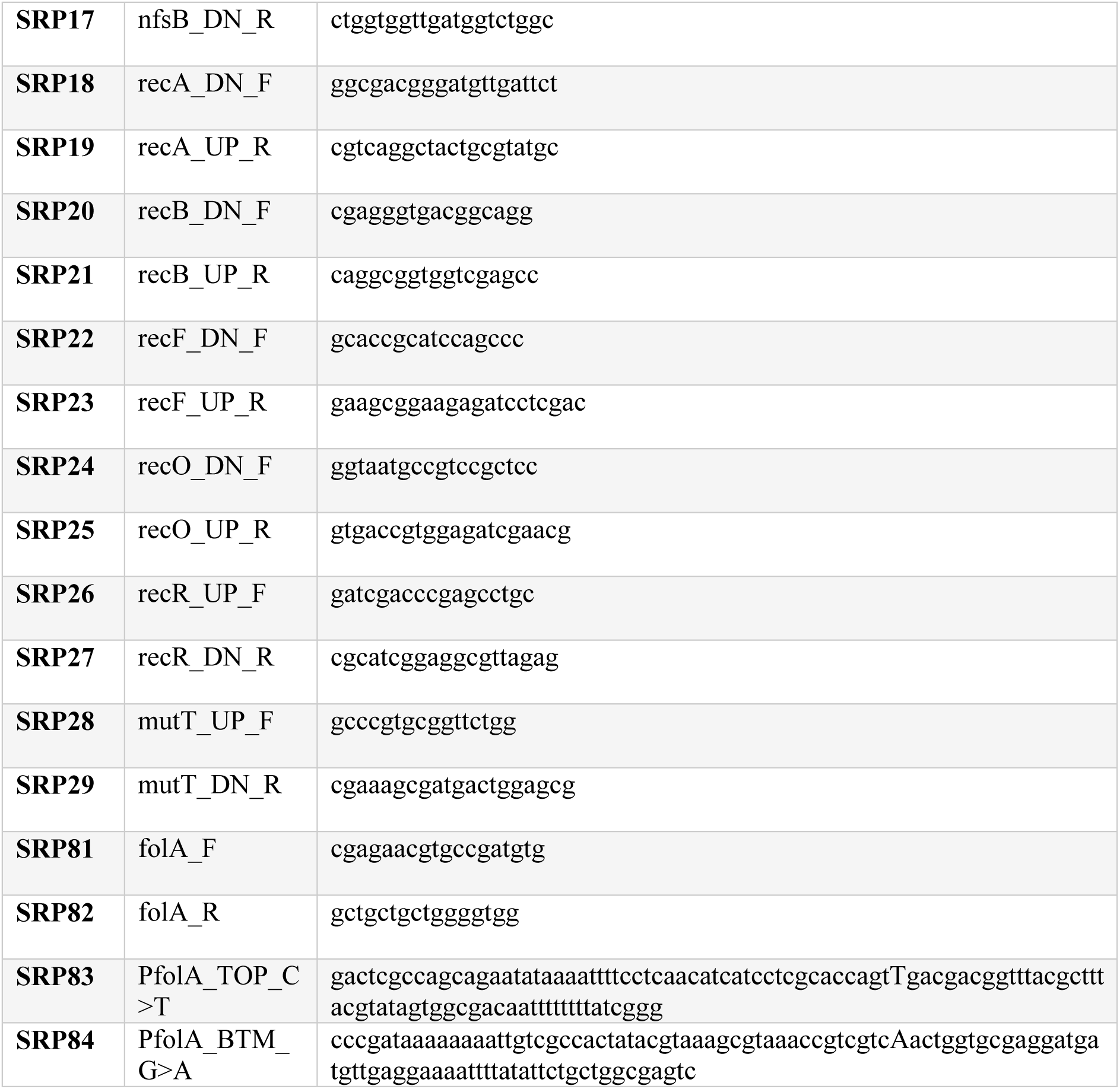
Oligonucleotides used in this study

### Strain construction and complementation

Mutant strains were constructed in the *E. coli* K12 MG1655 background (unless otherwise stated) using λ_RED_ recombination, replacing the gene of interest with a kanamycin cassette flanked by FRT sites ^84^. Construction of Δ*recA*::Kan^R^ (HH020) ^85^, Δ*recB*::Kan^R^ (EAW102) ^32^, Δ*recF*::Kan^R^ (EAW629), Δ*recO*::Kan^R^ (EAW114) and Δ*recR*::Kan^R^ (EAW669) ^86^ have been described previously. EAW999 Δ*mutT*::Kan^R^ was constructed via λ_RED_ recombination using pKD46 ^84^. Where applicable, the kanamycin resistance cassette was removed from strains via FLP-FRT recombination using the plasmid pLH29 ^87^ to obtain the kanamycin sensitive derivatives: HH021 (Δ*recA*::FRT), HG356 (Δ*recB*::FRT), MV009 (Δ*recF*::FRT), SRM019 (Δ*recO*::FRT), SRM020 (Δ*recR*::FRT) and MV005 (Δ*mutT*::FRT). All mutations were confirmed by PCR.

Mutant DNA repair alleles were moved into the ciprofloxacin-resistant (Cip^R^) background (CH5741) using P1 transduction. P1 phage lysates were raised using HH020 (Δ*recA*::Kan^R^), EAW102 (Δ*recB*::Kan^R^), EAW629 (Δ*recF*::Kan^R^), EAW114 (Δ*recO*::Kan^R^) and EAW669 (Δ*recR*::Kan^R^), EAW999 (Δ*mutT*::Kan^R^).

Kanamycin resistant (Kan^R^) strains were constructed by introducing the pUA66 plasmid (which confers kanamycin resistance through the *aph(3’)-*II gene) into the strains MG1655 (WT), HH021 (Δ*recA*::FRT), HG356 (Δ*recB*::FRT), MV009 (Δ*recF*::FRT), SRM019 (Δ*recO*::FRT), SRM020 (Δ*recR*::FRT) and MV005 (Δ*mutT*::FRT) via transformation.

Ampicillin resistant (Amp^R^) strains were constructed by introducing the pWSK29 plasmid (which confers ampicillin resistance through the *bla* gene) into the strains MG1655 (WT), HH020 (Δ*recA*::Kan^R^), EAW102 (Δ*recB*::Kan^R^), EAW629 (Δ*recF*::Kan^R^), EAW114 (Δ*recO*::Kan^R^). EAW669 (Δ*recR*::Kan^R^) and EAW999 (Δ*mutT*:: Kan^R^) via transformation.

Trimethoprim resistant (Tmp^R^) strains were constructed through λ RED recombination of SRP84 (Table 4) with MG1655. This recombination introduced a single point mutation from C > T in position 49765 of the chromosome (EKW048). Mutation was confirmed by PCR amplification and sequencing. P1 phage lysates were then raised using HH020 (Δ*recA*::Kan^R^), EAW102 (Δ*recB*::Kan^R^), EAW629 (Δ*recF*::Kan^R^), EAW114 (Δ*recO*::Kan^R^) and EAW669 (Δ*recR*::Kan^R^), EAW999 (Δ*mutT*::Kan^R^) and transduced with EKW048.

Nitrofurantoin resistant (Nit^R^) strains were constructed using P1 transduction. P1 phage lysate were raised on JW0835-1 (*nfsA*::kan^R^) from the Keio collection ^88^ and transduced into the strains MG1655 (WT), HH021 (Δ*recA*::FRT), HG356 (Δ*recB*::FRT), MV009 (Δ*recF*::FRT), SRM019 (Δ*recO*::FRT), SRM020 (Δ*recR*::FRT) and MV005 (Δ*mutT*::FRT). Kanamycin resistance was cured after each transduction and subsequently cured of the pLH29 plasmid to produce antibiotic sensitive variants.

For complementation of the Δ*recA* mutants, the plasmid, pHG134 (also referred to as pRecA), was used as described previously ^85^. For complementation of Δ*recB* mutants, the plasmid pSRM3 (referred to as pRecB) was constructed by Aldevron. The *recB* gene and 200 bp upstream was synthesised and cloned into KpnI/XbaI restriction sites on the pJM1071 plasmid backbone. Plasmid sequence is available in supplementary data. Complementation plasmids were introduced into the appropriate strains by transformation.

### Minimum inhibitory concentration (MIC) strip assays and TD tests

MICs were primarily determined using Liofilchem® MTS™ (MIC Test Strips) according to the manufacturers recommendations. When required, TD tests ^58^ were performed following the MIC strip assay. The antibiotic strip was removed from the plate and 5 µl of 40% (w/v) D-glucose solution was then added and left to dry at room temperature. Plates were further incubated overnight at 37°C. Tolerance was described as the growth of colonies in the zone of inhibition following the addition of glucose.

### Disc diffusion assays and TD tests

Where MIC test strips were unavailable or a strains MICs were beyond the strip test range, disc diffusion assays were performed. Cells were grown in 500 µl LB broth for roughly 6 hours at 37°C. Then, 100 µl of culture was plated onto a LB agar plate. A sterile 13 mm Whatman® disc (GE Healthcare) with antibiotic (or compound) was placed in the centre of the plate. Final antibiotic/ compound concentrations on the discs were as follows unless otherwise stated: 10 mg ampicillin, 1 mg trimethoprim, 3.5 mg kanamycin, 50 µg ML328, 15 µg IMP-1700. Plates were incubated 37°C overnight. The zone of inhibition surrounding the disc was measured. When required, a modified TD test ^58^ was performed following the disc diffusion assay as described above.

### Tolerance regrowth percentage and zone of inhibition area calculations

Regrowth percentages and the area within the zone of inhibition (ZOI) were measured using ImageJ ^89^. Areas, and subsequent regrowth, under the MIC strip or antibiotic disc were excluded from these measurements. Images of MIC plates were imported, cropped and aligned using regular ImageJ functions. To select the ZOI, images were subject to thresholding using the “auto-threshold” mean pre-set function. The ZOI was selected using the “wand” tool and added to the ROI manager. Scale was set to pixels and area was measured using the “measurements function”.

For TD image processing, ZOI selection and measurements were conducted as above. To select for any colonies within the ZOI, all thresholded regions were selected using the “select all” function and added to the ROI manager. Any bacterial growth already present within the ZOI were included in measurments as follows. Area within the ZOI without bacterial growth was determined by selecting the two ROIs and using the “AND” and measurement functions. TD images were cropped and aligned as above. TD images were subject to thresholding using the “auto-threshold” mean pre-set function. The ZOI from the MIC plates was added to the TD image and position adjusted as needed. Colonies were selected as before using the “select all” function. Area within the ZOI without colonies were measured using the “AND” and measurement functions. Measurements were exported and further processed in excel. Percent regrowth was calculated using the following equation:

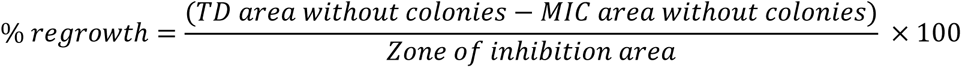

Any area or regrowth under antibiotic strips or discs were not included in the re-growth measurements. The ImageJ macro code used to analyse images is available in the supplementary data.

### Time-kill experiments

To determine the viability of wild-type and mutant strains, viability was examined using time-kill assays. Single colonies were used to inoculate 1 ml Mueller-Hinton (MH) cation adjusted broth and incubated 850 rpm at 37°C overnight. The overnight culture was reset 1:100 in 10 ml MH broth and incubated shaking 200 rpm for until an OD_600_ of 0.4 was reached. Where needed, OD_600_ values were normalised to 0.4. A 2 ml sample of culture was then incubated with 3 µg/ml (3 X MIC), 5 µg/ml (5 X MIC), or 10 µg/ml (10 X MIC) kanamycin for three hours. Samples (200 µl) were taken hourly and washed three times in 0.1M MgSO_4_. Serial dilutions were prepared in LB to 10^-6^ and 5 µl of each dilution was spot in duplicate onto MH agar plates. Plates were dried and incubated at 37°C overnight. Colonies were enumerated and cfu/ml determined for each time-point.

### Antibiotic sensitivity assays (Spot-plate assays)

Single colonies were used to inoculate 1 ml LB broth and incubated 850 rpm at 37°C overnight. The following day, overnight culture was reset 1:100 in LB broth and incubated shaking until an OD_600_ of 0.2 was reached. Serial dilutions were prepared to 10^-5^ and 5 µl of each dilution was spot, in duplicate, onto LB agar plates and LB agar with antibiotic. Antibiotics were added to final concentrations of: 650 and 775 µg/ml kanamycin, 0.2 µg/ml trimethoprim (sensitive), 5 µg/ml trimethoprim (Tmp^R^). Plates were allowed to dry and incubated at 37°C overnight. Plates were imaged using a BioRad GelDoc imager.

### MIC/minimum bactericidal concentration (MBC) broth assays

Where necessary, MICs were determined using a 96-well plate in accordance with the EUCAST broth microdilution protocol. Single colonies were used to inoculate 1 ml Mueller-Hinton (MH) cation adjusted broth and incubated 37°C overnight. The overnight culture was diluted 1:100 in MH broth and incubated shaking 850 rpm for three hours. Exponential phase cells were then diluted 1:200. 10 µl of culture (approx. 10^5^ cfu/mL) was used to inoculate wells containing 190 µl MH broth supplemented with DMSO/compound, ensuring a 2% (v/v) DMSO final concentration in all wells. Compounds were prepared in two-fold serial dilutions. Plates were incubated at 37°C in PolarStar Omega plate reader with shaking. The optical density at 600 nm (OD_600_) was recorded every 20 minutes for 18 hours. OD_600_ measurements were background corrected against no-inoculum controls. The MIC was defined as the lowest concentration of compound with no growth as determined by OD_600_ readings. MIC values were calculated using data from at least three biological replicates. IC_50_ values were determined by dose-response non-linear regression.

### SOS plate assays

SOS induction in response to antibiotic exposure was analysed qualitatively by an agar plate based fluorescence assay. This assay used strains containing a reporter plasmid that expresses the fast folding GFP derivative, GFPmut2 ^48^, under the control of the SOS-inducible promoter P*_sulA._* Cells were grown in 500 µl LB broth supplemented with kanamycin (to select for the reporter plasmid) for roughly 6 hours at 37°C. Then, 100 µl of culture was used to inoculate 4 ml of soft LB agar (0.5% agar (w/v)).

The agar was then poured on top of a regular LB Kan plate (to maintain selection for the reporter plasmid) and allowed to set. A Liofilchem® MTS™ (MIC Test Strips), or sterile 13 mm Whatman® disc (GE Healthcare) with antibiotic was placed in the centre of the plate. Plates were incubated 37°C for 24 hours. Fluorescence signal was detected using a custom-built fluorescence photography set up using an Andor Zyla 5.5 camera equipped with a Nikon 18-55 mm SLR objective and a Chroma ZET405/488/594m-TRF emission filter. Excitation was achieved with a Thorlabs DC4104 high power LED source behind a Chroma ZET405/488/594x excitation filter. GFP was visualised using 490 nm wavelength excitation at a power of 350 mW. Exposure times were adjusted as needed according to signal strength. The camera and LED light source were controlled using MicroManager software ^90^.

## Data availability

The data from this study are available from the corresponding author upon request.

