## Supplementary Data for "Defects in DNA double-strand break repair re-sensitise antibiotic-resistant *Escherichia coli* to multiple bactericidal antibiotics"

#### Supplementary Figure

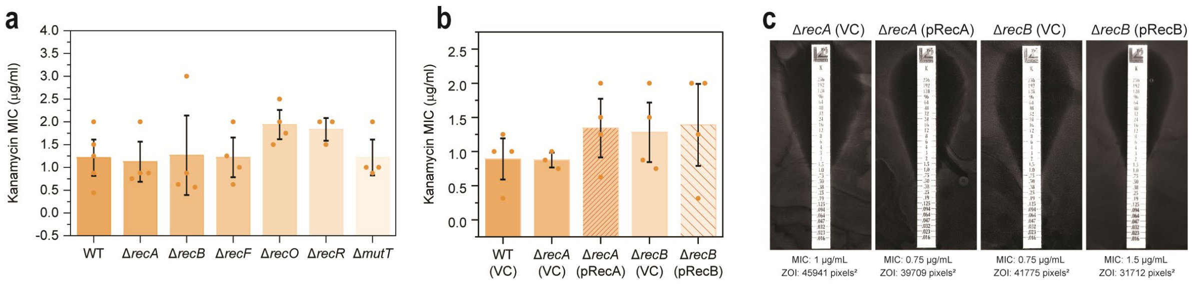


**Supplementary Figure S1: a.** Kanamycin MIC values obtained for isogenic *E. coli* strains MG1655 (WT; wild-type), Δ*recA*::Kan$^R$ (HH020), Δ*recB*::Kan$^R$ (EAW102), Δ*recF*::Kan$^R$ (EAW629), Δ*recO*::Kan$^R$ (EAW114), Δ*recR*::Kan$^R$ (EAW669) and Δ*mutT*::Kan$^R$ (EAW999). MICs were assayed using MIC test strips according to manufacturer's instruction. The means and standard errors of the mean are shown, based on results from at least four biological replicates. **b.** Kanamycin MIC values obtained for wild-type (MG1655) with empty vector (VC; vector control), Δ*recA* and Δ*recB* mutants with empty vector and complemented derivatives (pRecA and pRecB, respectively). The means and standard errors of the mean are shown based on results from at least three biological replicates. **c.** Representative images of kanamycin MIC plate assays for Δ*recA* (VC), Δ*recA* (pRecA), Δ*recB* (VC) and Δ*recB* (pRecB) *E. coli* strains. MICs and measured zone of inhibition areas in pixels$^2$ are denoted below the corresponding images.

### Supplementary Figure

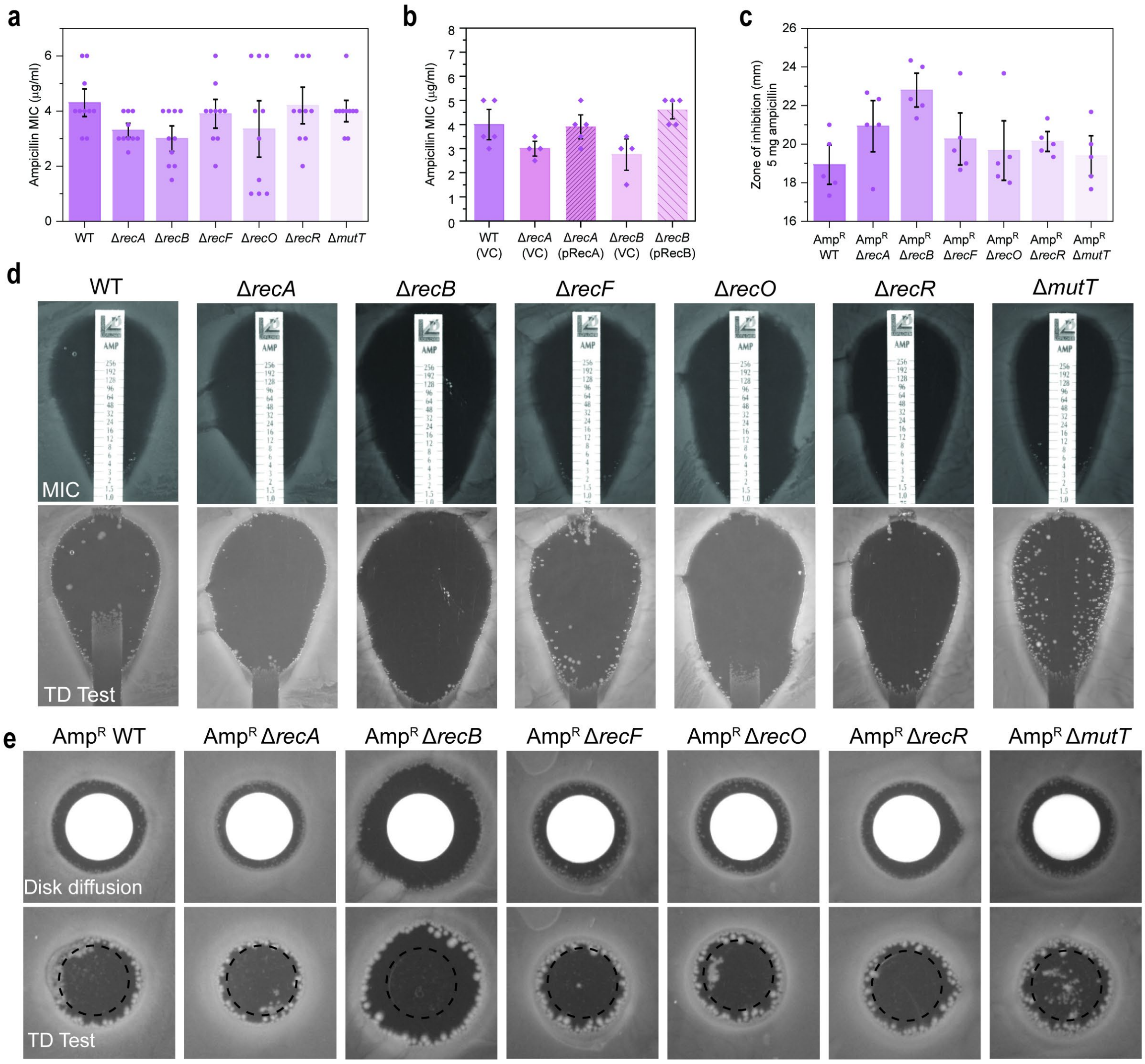


**Supplementary Figure S2: a.** Ampicillin MIC values obtained for isogenic *E. coli* strains MG1655 (WT; wild-type), Δ*recA*::Kan$^R$ (HH020), Δ*recB*::Kan$^R$ (EAW102), Δ*recF*::Kan$^R$ (EAW629), Δ*recO*::Kan$^R$ (EAW114), Δ*recR*::Kan$^R$ (EAW669) and Δ*mutT*::Kan$^R$ (EAW999). MICs were assayed using MIC test strips according to manufacturer's instruction. The means and standard errors of the mean are shown, based on results from at least three biological replicates. **b.** Ampicillin MIC values obtained for the *E. coli* strains wild-type (MG1655) with empty vector (VC; vector control), Δ*recA* and Δ*recB* mutants with empty vector and complemented derivatives (pRecA and pRecB, respectively). The means and standard errors of the mean are shown based on results from at least three biological replicates. Statistical analysis was carried out using a student's *t*-test. An asterisk denotes statistical significance ($p < 0.05$) compared to wild-type with empty vector, WT (VC). **c.** Zone of inhibition area measurements for ampicillin-resistant WT and DNA repair mutant strains following disk diffusion assays with 5 mg ampicillin. $Amp^R$ (COF001), $Amp^R$ Δ*recA* (COF002), $Amp^R$ Δ*recB* (COF003), $Amp^R$ Δ*recF* (COF004), $Amp^R$ Δ*recO* (COF005), $Amp^R$ Δ*recR* (COF006) and $Amp^R$ Δ*mutT* (COF007). The means and standard errors of the mean are shown based on results from at least four biological replicates. **d.** Representative images of ampicillin MIC and tolerance (TD Test) plate assays for isogenic *E. coli* strains MG1655 (WT; wild-type), Δ*recA*::$Kan^R$ (HH020), Δ*recB*::$Kan^R$ (EAW102), Δ*recF*::$Kan^R$ (EAW629), Δ*recO*::$Kan^R$ (EAW114), Δ*recR*::$Kan^R$ (EAW669) and Δ*mutT*::$Kan^R$ (EAW999). Images show representative plates from independent triplicate replicates. **e.** Representative images of ampicillin MIC and tolerance (TD Test) plate assays for isogenic *E. coli* strains $Amp^R$ (COF001), $Amp^R$ Δ*recA* (COF002), $Amp^R$ Δ*recB* (COF003), $Amp^R$ Δ*recF* (COF004), $Amp^R$ Δ*recO* (COF005), $Amp^R$ Δ*recR* (COF006) and $Amp^R$ Δ*mutT* (COF007). Images show representative plates from independent triplicate replicates.

**Supplementary Figure**

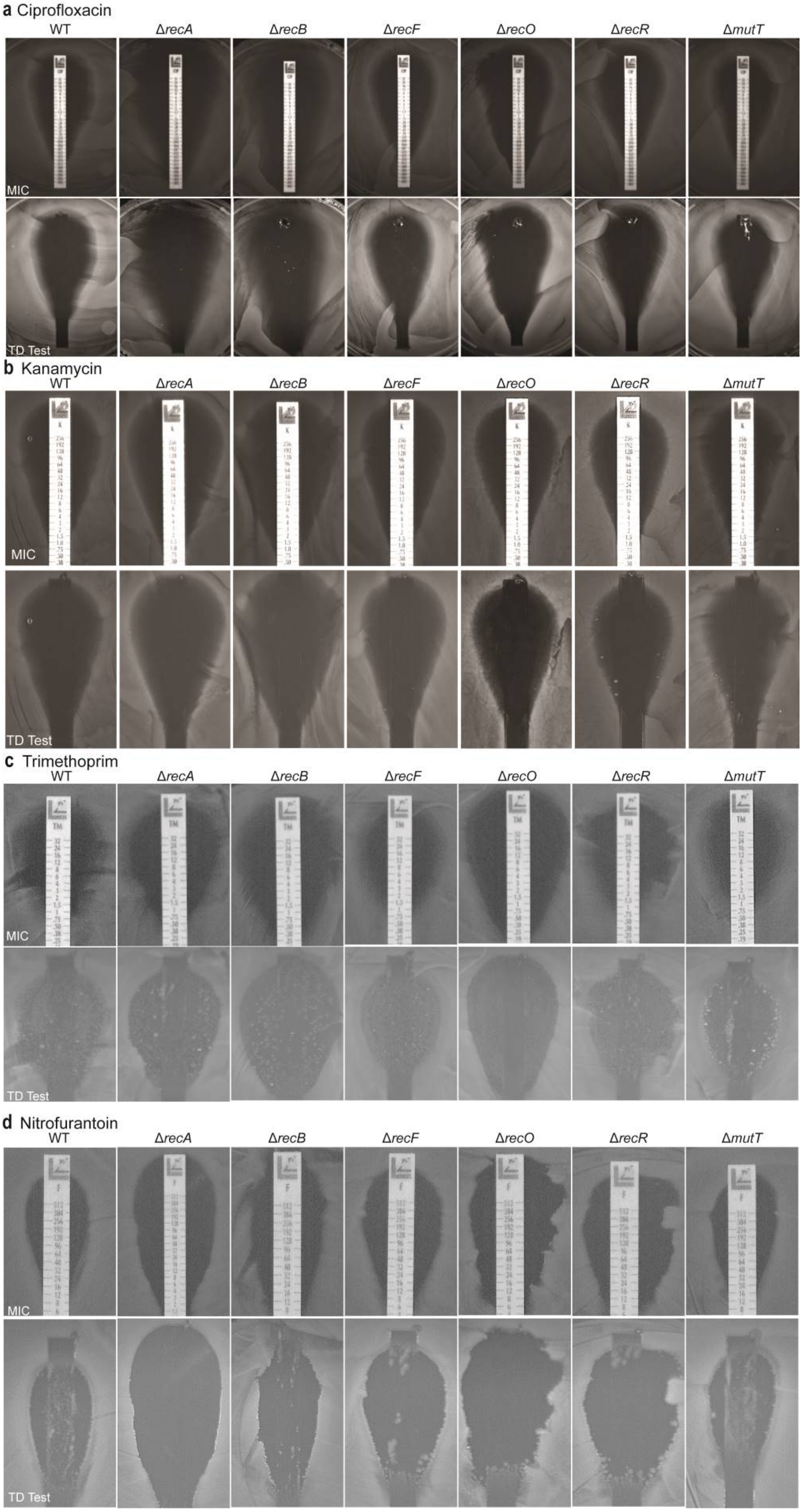

**Supplementary Figure S3:** **a.** Representative images of ciprofloxacin MIC and tolerance (TD Test) plate assays for wild-type (WT), and DNA repair deficient *E. coli* strains. **b.** Representative images of kanamycin MIC and tolerance (TD Test) plate assays for wild-type (WT), and DNA repair deficient *E. coli* strains. **c.** Representative images of nitrofurantoin MIC and tolerance (TD Test) plate assays for wild-type (WT), and DNA repair deficient *E. coli* strains. **d.** Representative images of trimethoprim MIC and tolerance (TD Test) plate assays for wild-type (WT), and DNA repair deficient *E. coli* strains.

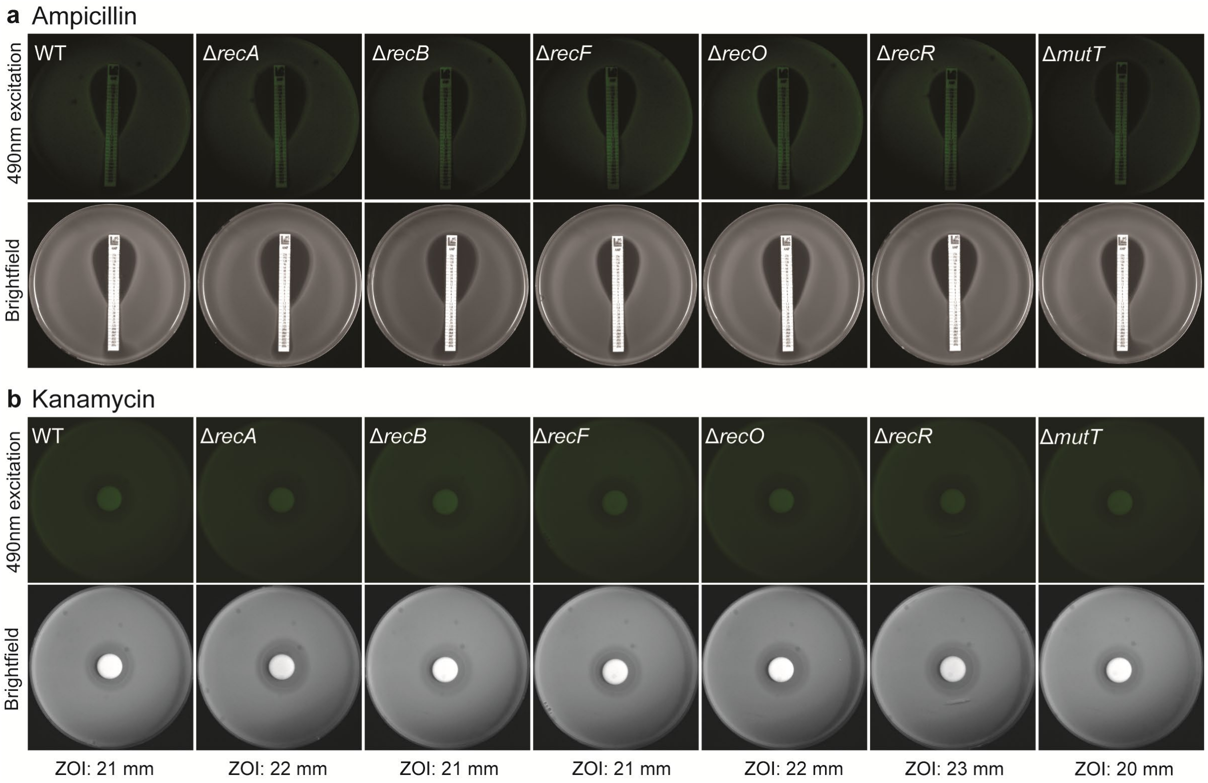


**Supplementary Figure S4: a.** Expression of SOS reporter fusion $P_{sulA}$-*gfp* on a solid agar surface in wild-type (WT) and DNA repair deficient strains grown in the presence of ampicillin (0.016 - 256 µg/ml). Plates were visualised under 490 nm excitation (top panels) and in bright-field (lower panel). Any SOS induction should be visualised as strong fluorescence band at the border of the zone of inhibition. **b.** Expression of SOS reporter fusion $P_{recN}$-*gfp* on a solid agar surface in wild-type (WT) and DNA repair deficient strains grown in the presence of 50 µg kanamycin. Plates were visualised under a 490 nm excitation (top panels) and in bright-field (lower panel). The zone of inhibition recorded for each strain is denoted below the bright-field image.

### Supplementary Fig.

**a** ML328 MIC $OD_{600}$ and $IC_{50}$ linear-regession

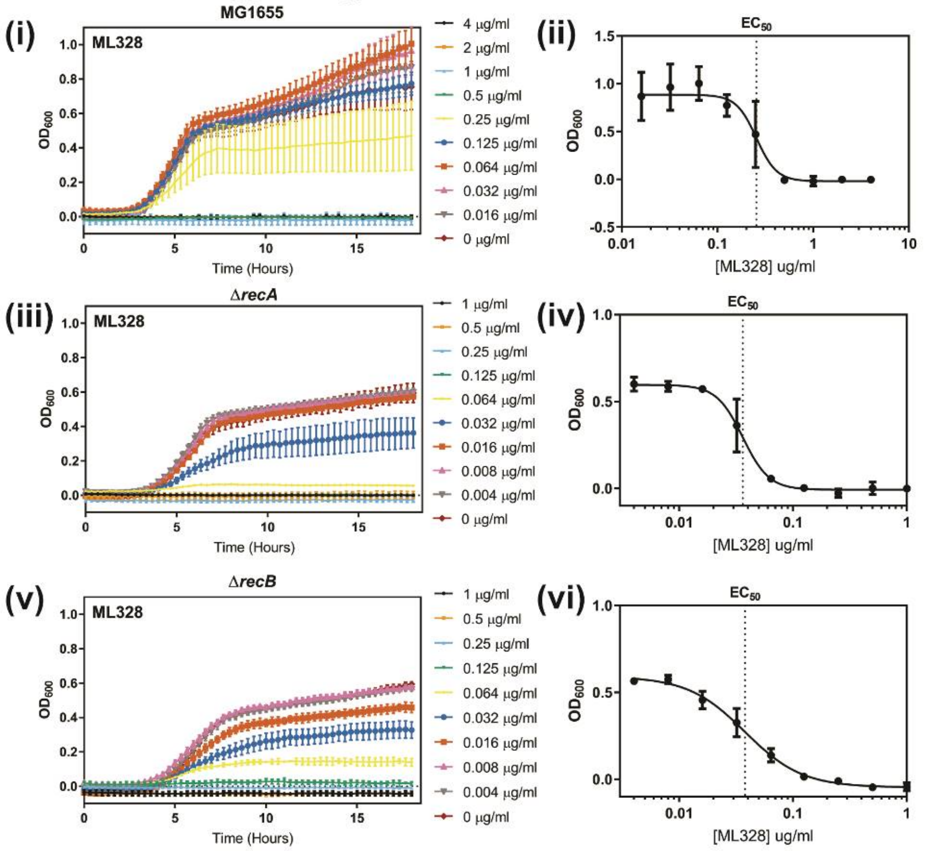


**b** IMP-1700 MIC $OD_{600}$ and $IC_{50}$ linear-regession

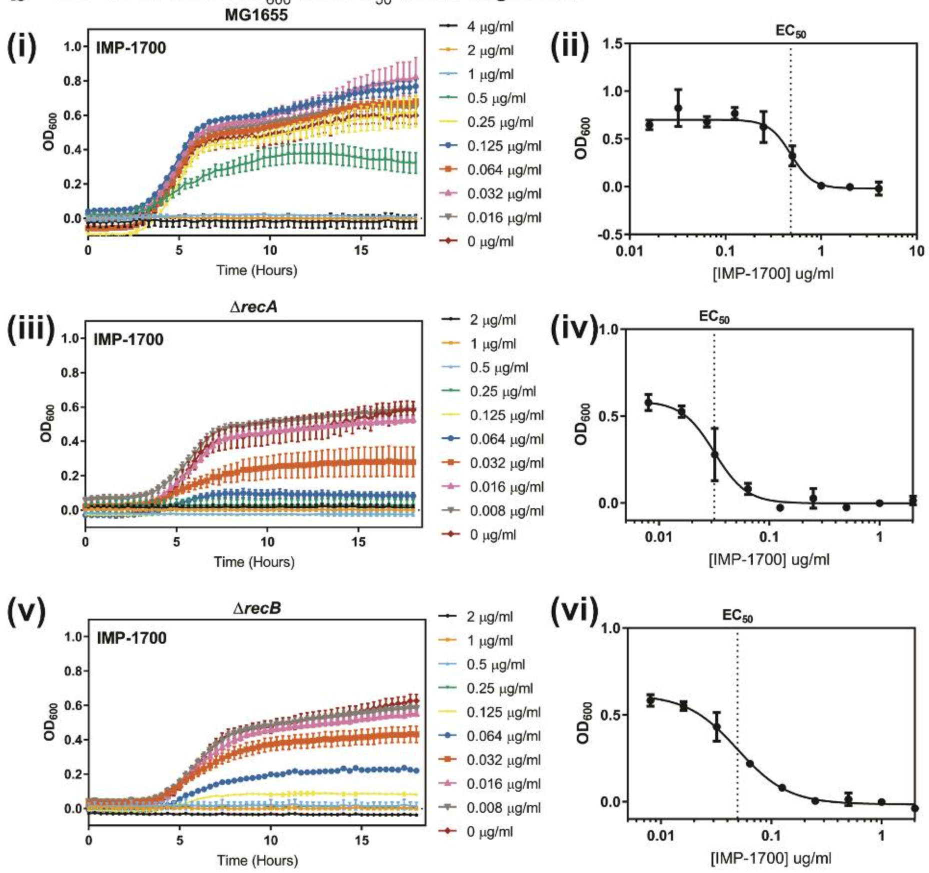

**Supplementary Figure S5**: ML328 **(a.)** and IMP-1700 **(b.)** $OD_{600}$ and $IC_{50}$ linear regression data for *E. coli* strains (**i** and **ii**) MG1655 (WT; wild-type), (**iii** and **iv**) Δ*recA*::$Kan^R$ (HH020), (**v** and **vi**) Δ*recB*::$Kan^R$ (EAW102). The optical density at 600 nm ($OD_{600}$) was recorded every 20 minutes for 18 hours. $OD_{600}$ measurements were background corrected against no-inoculum controls. Shown are the means and standard error of the mean from three biological replicates. The MIC was defined as the lowest concentration of compound with no growth as determined by $OD_{600}$ readings. $IC_{50}$ values were calculated using data from at least three biological replicates.

### Supplementary Figure

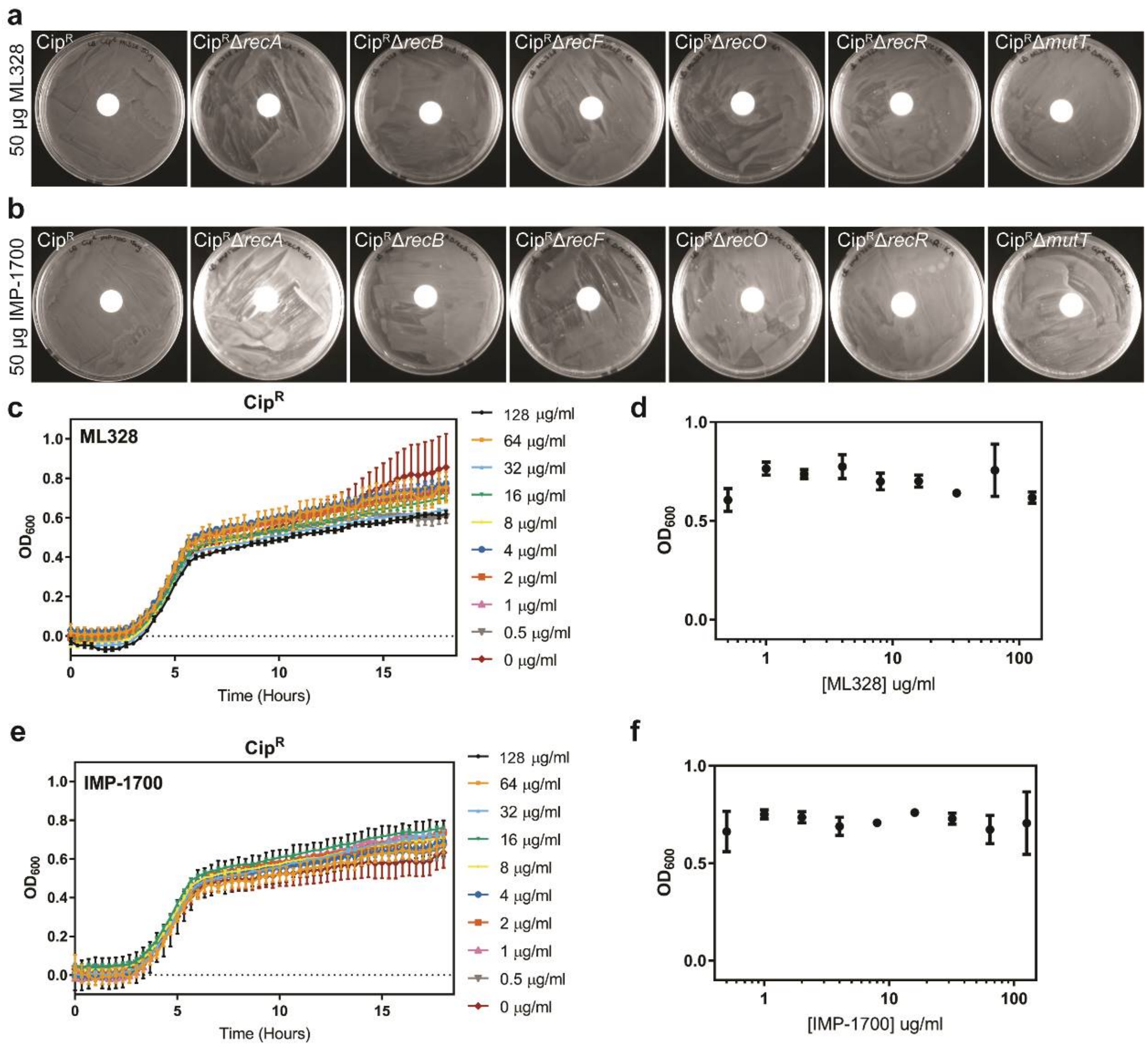


**Supplementary Figure S6**: **a. and b.** Representative images of ML328 (**a.**) and IMP-1700 (**b.**) disk diffusion assays obtained for isogenic ciprofloxacin-resistant ($Cip^R$) DNA repair deficient *E. coli* strains $Cip^R$ (CH5741), $Cip^R$ Δ*recA* (FM002), $Cip^R$ Δ*recB* (FM001), $Cip^R$ Δ*recF* (FM003), $Cip^R$ Δ*recO* (FM004), $Cip^R$ Δ*recR* (FM005) and $Cip^R$ Δ*mutT* (MV001). ML328 (**c. and d.**) and IMP-1700 (**e. and f.**) $OD_{600}$ and $IC_{50}$ linear regression data for *E. coli* strains. The optical density at 600 nm ($OD_{600}$) was recorded every 20 minutes for 18 hours. $OD_{600}$ measurements were background corrected against no-inoculum controls. Shown are the means and standard error of the mean from three biological replicates. $IC_{50}$ values were calculated using data from at least three biological replicates.

**Supplementary Movie 1:** Time-lapse acquisition (5 minute intervals) of the expression of SOS reporter fusion $P_{sulA}$-*gfp* on a solid agar surface in wild-type (WT) *E. coli* grown in the presence of a ciprofloxacin MIC test strip (0.002-32 µg/ml). SOS induction is visualised as strong fluorescence band at the border of the zone of inhibition.

**Supplementary data:** ImageJ Macros to analyse percentage regrowth

**IJ1MACRO** 1: (measuring MIC plate ZOI area and area without colonies)

```
//step 1: duplicate image and normalise brightness and contrast
//NOTE: Renaming only works if the ROI manager is empty
// clears results table and ROI manager
run("Clear Results");
selectWindow("ROI Manager");
run("Close");
//run("Brightness/Contrast...");
run("Duplicate...", " ");
setMinAndMax(0, 4079);
//step 2: threshold the zone of inhibition using a preset auto-
threshold
setAutoThreshold("Default no-reset");
setAutoThreshold("Mean stack");
// step 3: select the zones of inhibition and add to ROI manager
//setTool("wand");
doWand(496, 300);
roiManager("Add");
doWand(400, 300);
roiManager("Add");
roiManager("Select", newArray(0,1));
roiManager("Combine");
roiManager("Add");
roiManager("Select", 2);
roiManager("Rename", "ZOI");
//step 4: select all thresholded regions (this includes colonies in
ZOI)
run("Create Selection");
roiManager("Add");
roiManager("Select", 3);
roiManager("Rename", "Colonies");
//step 5: set scale to pixels and measure zone of inhibition
run("Set Scale...", "distance=0 known=0 pixel=1 unit=pixel global");
roiManager("Select", 2);
run("Measure");
//step 6: measure growth in ZOI
roiManager("Select", newArray(2,3));
roiManager("AND");
run("Measure");
```

IJ1MACRO 2: (measuring the area without colonies on the TD plate)

```
//step 1: duplicate image and normalise brightness and contrast
//NOTE: Renaming only works if the ROI manager is empty
//run("Brightness/Contrast...");
run("Duplicate...", " ");
setMinAndMax(0, 4079);
//step 2: threshold the zone of inhibition using a preset auto-
threshold
setAutoThreshold("Default no-reset");
setAutoThreshold("Mean stack");
// step 3: add the original zone of inhibition to ROI manager
```

```
roiManager("Select", 2);
waitForUser("Move ROI to an appropriate position")
wait(3000)
roiManager("Add");
roiManager("Select", 4);
roiManager("Rename", "ZOI-TD");
//step 4: select all thresholded regions (this includes colonies in
ZOI)
run("Create Selection");
roiManager("Add");
roiManager("Select", 5);
roiManager("Rename", "Colonies-TD");
//step 5: set scale to pixels and measure area of colonies in zone of
inhibition
run("Set Scale...", "distance=0 known=0 pixel=1 unit=pixel global");
roiManager("Select", newArray(4,5));
roiManager("AND");
run("Measure");
//step 6: save results as csv
selectWindow("Results");
saveAs("Results")
selectWindow("ROI Manager");
```

**Supplementary Data:** Sequence of pSRM3 (pRecB complementation plasmid)

AAGCTGGAAGATCTTCCCTGGCACGACAGGTTTCCCGACTGGAAAGCGGGCAGTGAGCGCAACGCAAT
TAATGTGAGTTAGCTCACTCATTAGGCACCCCAGGCTTTACACTTTATGCTTCCGGCTCGTATGTTGT
GTGGAATTGTGAGCGGATAACAATTTCACACAGGAAACAGCTATGACCATGATTACGCCAAGCGCGCA
ATTAACCCTCACTAAAGGGAACAAAAGCTGGGTACCGGCCCAGATAAAACTGCTGACGCCGCAAAAAC
TTGCTGATTTCTTCCATCAGGCGGTGGTCGAGCCGCAAGGCATGGCTATTCTGTCGCAGATTTCCGGC
AGCCAGAACGGGAAAGCCGAATATGTACACCCTGAAGGCTGGAAAGTGTGGGAGAACGTCAGCGCGTT
GCAGCAAACAATGCCCCTGATGAGTGAAAAGAATGAGTGATGTCGCCGAGACACTAGATCCTTTGCGC
TTGCCCTTACAGGGTGAGCGCCTGATTGAAGCCTCTGCCGGCACAGGCAAAACCTTTACGATTGCGGC
GCTCTATTTGCGCCTGTTACTTGGACTAGGCGGTTCCGCCGCCTTTCCCCGCCCGCTGACCGTTGAAG
AACTGCTGGTGGTCACCTTTACCGAGGCTGCCACGGCAGAATTGCGCGGTCGTATCCGTAGCAATATC
CACGAGTTGCGCATCGCCTGTCTGCGTGAAACCACCGACAATCCACTGTACGAACGCCTGCTGGAAGA
GATCGACGATAAAGCGCAAGCCGCGCAGTGGTTGTTGTTAGCCGAACGGCAGATGGATGAAGCGGCAG
TCTTTACTATTCACGGCTTTTGCCAGCGCATGCTCAACCTGAATGCCTTTGAATCCGGCATGCTGTTT
GAGCAGCAGCTGATTGAAGATGAGTCTCTGCTACGCTACCAGGCCTGCGCCGATTTCTGGCGTCGCCA
CTGCTACCCGCTGCCGCGTGAAATAGCCCAGGTCGTCTTTGAAACCTGGAAAGGGCCGCAGGCGTTGC
TGCGCGATATTAATCGTTATCTGCAAGGCGAAGCGCCGGTTATCAAAGCACCGCCGCCCGATGATGAA
ACGCTGGCTTCCCGTCACGCGCAAATTGTGGCGCGTATTGATACGGTAAAACAGCAGTGGCGCGACGC
AGTGGGTGAACTGGATGCGCTGATCGAATCTTCTGGTATTGATCGACGCAAGTTTAACCGTAGCAATC
AGGCTAAATGGATCGACAAGATCAGCGCCTGGGCAGAAGAAGAGACAAACAGTTATCAGTTGCCGGAG
TCGCTGGAAAAATTCTCCCAGCGTTTCTTAGAAGATCGCACGAAGGCCGGGGGGGAAACCCCGCGACA
TCCACTGTTTGAGGCGATCGATCAACTGCTTGCAGAACCATTGTCGATCCGCGATCTGGTGATCACCC
GCGCATTGGCTGAGATCCGCGAAACAGTAGCGCGTGAAAAACGCCGCCGTGGCGAATTGGGTTTTGAT
GACATGTTAAGTCGGCTCGATTCCGCGCTGCGTAGCGAAAGCGGTGAGGTGTTGGCAGCGGCGATCCG
TACGCGATTCCCGGTGGCAATGATCGATGAATTTCAGGATACCGACCCCCAGCAGTACCGAATTTTTC
GCCGTATCTGGCACCATCAGCCGGAAACCGCATTGTTGCTAATTGGCGACCCGAAGCAGGCCATATAT
GCATTCCGGGGTGCGGATATCTTCACTTATATGAAGGCGCGTAGCGAAGTTCACGCCCACTACACTTT
AGACACCAACTGGCGTTCCGCACCAGGAATGGTGAACAGCGTGAATAAGCTTTTCAGCCAGACTGATG
ACGCGTTCATGTTTCGCGAAATACCGTTTATTCCAGTGAAATCAGCCGGGAAAAATCAGGCGTTACGT
TTTGTATTTAAAGGTGAAACACAGCCTGCGATGAAAATGTGGCTGATGGAAGGCGAAAGCTGCGGCGT
TGGCGATTATCAAAGTACCATGGCGCAGGTATGTGCTGCGCAAATCCGCGACTGGCTACAAGCCGGAC
AGCGGGGCGAAGCGTTGCTGATGAACGGCGACGACGCGCGTCCGGTGCGTGCTTCGGACATCAGTGTG
CTGGTGCGCAGCCGCCAGGAGGCCGCCCAGGTGCGCGATGCCTTAACGTTGCTGGAAATCCCTTCCGT
TTACCTTTCGAACCGCGACAGTGTTTTTGAAACTCTGGAAGCGCAGGAAATGCTTTGGTTGTTGCAGG
CGGTGATGACGCCCGAACGTGAGAACACCCTGCGTAGTGCGCTGGCAACGTCAATGATGGGGCTGAAC
GCGCTGGATATCGAAACGCTGAACAATGACGAACATGCGTGGGATGTGGTAGTCGAAGAGTTCGATGG
TTATCGGCAAATCTGGCGCAAACGTGGCGTTATGCCGATGCTGCGGGCGCTGATGTCGGCGCGTAACA
TTGCTGAAAACTTGCTGGCAACGGCAGGCGGTGAGCGGCGTCTTACCGATATCTTGCATATCAGCGAA
CTGCTACAAGAAGCCGGAACGCAGCTGGAAAGTGAACATGCGCTGGTACGCTGGTTATCGCAACATAT
CCTCGAGCCAGACAGTAATGCCTCCAGCCAACAAATGCGTCTCGAAAGTGATAAACATCTGGTGCAGA
TTGTCACGATCCACAAATCGAAAGGGCTGGAATATCCATTGGTCTGGCTGCCGTTTATCACCAATTTC
CGCGTCCAGGAGCAGGCGTTTTATCACGATCGCCACTCGTTTGAGGCAGTTCTGGATCTTAATGCTGC
GCCAGAAAGCGTCGACCTCGCGGAGGCCGAACGTCTGGCGGAAGATCTGCGTTTGCTTTACGTGGCGC
TGACACGTTCGGTTTGGCATTGCAGTCTCGGCGTTGCACCGCTGGTGCGCCGTCGTGGCGATAAAAAA
GGTGACACCGACGTCCACCAAAGTGCGCTCGGGCGTTTGCTGCAAAAAGGGGAACCGCAAGATGCGGC
AGGGCTTCGCACCTGTATTGAAGCGTTATGCGATGATGATATTGCCTGGCAAACGGCACAAACTGGTG
ATAACCAACCCTGGCAGGTTAATGATGTTTCTACAGCAGAGCTGAATGCGAAGACGTTACAACGATTG
CCCGGCGATAACTGGCGCGTCACCAGCTACTCTGGTTTGCAACAGCGTGGTCACGGTATCGCCCAGGA
TTTGATGCCTCGGCTGGATGTCGATGCTGCAGGCGTTGCCAGCGTCGTTGAAGAACCGACGTTAACAC
CACATCAGTTTCCGCGCGGTGCGTCACCGGGGACGTTCTTGCACAGTTTGTTTGAAGACCTGGATTTT
ACCCAGCCGGTTGACCCGAACTGGGTGCGGGAAAAACTGGAACTCGGCGGCTTTGAATCGCAGTGGGA
ACCGGTATTGACCGAGTGGATCACGGCTGTCCTCCAGGCACCTCTCAATGAAACCGGCGTAAGCCTGA
GTCAACTTTCCGCCCGCAATAAACAGGTGGAGATGGAGTTTTATCTGCCGATTAGTGAACCGCTTATC
GCCAGTCAGCTTGATACGTTAATCCGCCAGTTTGACCCGCTATCCGCAGGCTGCCCGCCGCTGGAGTT

CATGCAGGTACGTGGCATGTTAAAAGGCTTTATCGACCTGGTGTTCCGCCACGAAGGGCGTTATTACC
TGCTCGACTATAAATCCAACTGGTTGGGTGAAGACAGTTCGGCTTACACCCAACAGGCTATGGCAGCG
GCAATGCAGGCACACCGCTATGATCTGCAATATCAGCTTTATACCCTGGCGCTGCATCGTTATCTGCG
CCATCGCATTGCTGATTACGACTATGAGCACCACTTTGGCGGCGTTATTTATCTGTTCCTGCGTGGCG
TTGATAAAGAACATCCGCAACAGGGGATTTACACAACCCGACCCAACGCCGGGTTGATTGCCCTGATG
GATGAGATGTTTGCCGGTATGACCCTGGAGGAGGCGTAATCTAGAGCGGCCGCCACCGCGGTGGAGCT
CCAATTCGCCCTATAGTGAGTCGTATTACGCGCGCTCACTGGCCGTCGTTTTACAACGTCGTGACTGG
GAAAACCCTGGCGTTACCCAACTTAATCGCCTTGCAGCACATCCCCCTTTCGCCAGCTGGCGTAATAG
CGAAGAGGCCCGCACCGATCGCCCTTCCCAACAGTTGCGCAGCCTGAATGGCGAATGGGACGCGCCCT
GTAGCGGCGCATTAAGCGCGGCGGGTGTGGTGGTTACGCGCAGCGTGACCGCTACACTTGCCAGCGCC
CTAGCGCCCGCTCCTTTCGCTTTCTTCCCTTCCTTTCTCGCCACGTTCGCCGGAAGATCTTCCAATTC
CCGACAGTAAGACGGGTAAGCCTGTTGATGATACCGCTGCCTTACTGGGTGCATTAGCCAGTCTGAAT
GACCTGTCACGGGATAATCCGAAGTGGTCAGACTGGAAAATCAGAGGGCAGGAACTGCTGAACAGCAA
AAAGTCAGATAGCACCACATAGCAGACCCGCCATAAAACGCCCTGAGAAGCCCGTGACGGGCTTTTCT
TGTATTATGGGTAGTTTCCTTGCATGAATCCATAAAAGGCGCCTGTAGTGCCATTTACCCCCATTCAC
TGCCAGAGCCGTGAGCGCAGCGAACTGAATGTCACGAAAAAGACAGCGACTCAGGTGCCTGATGGTCG
GAGACAAAAGGAATATTCAGCGATTTGCCCGAGCTTGCGAGGGTGCTACTTAAGCCTTTAGGGTTTTA
AGGTCTGTTTTGTAGAGGAGCAAACAGCGTTTGCGACATCCTTTTGTAATACTGCGGAACTGACTAAA
GTAGTGAGTTATACACAGGGCTGGGATCTATTCTTTTTATCTTTTTTTATTCTTTCTTTATTCTATAA
ATTATAACCACTTGAATATAAACAAAAAAAACACACAAAGGTCTAGCGGAATTTACAGAGGGTCTAGC
AGAATTTACAAGTTTTCCAGCAAAGGTCTAGCAGAATTTACAGATACCCACAACTCAAAGGAAAAGGA
CTAGTAATTATCATTGACTAGCCCATCTCAATTGGTATAGTGATTAAAATCACCTAGACCAATTGAGA
TGTATGTCTGAATTAGTTGTTTTCAAAGCAAATGAACTAGCGATTAGTCGCTATGACTTAACGGAGCA
TGAAACCAAGCTAATTTTATGCTGTGTGGCACTACTCAACCCCACGATTGAAAACCCTACAAGGAAAG
AACGGACGGTATCGTTCACTTATAACCAATACGCTCAGATGATGAACATCAGTAGGGAAAATGCTTAT
GGTGTATTAGCTAAAGCAACCAGAGAGCTGATGACGAGAACTGTGGAAATCAGGAATCCTTTGGTTAA
AGGCTTTGAGATTTTCCAGTGGACAAACTATGCCAAGTTCTCAAGCGAAAAATTAGAATTAGTTTTTA
GTGAAGAGATATTGCCTTATCTTTTCCAGTTAAAAAAATTCATAAAATATAATCTGGAACATGTTAAG
TCTTTTGAAAACAAATACTCTATGAGGATTTATGAGTGGTTATTAAAAGAACTAACACAAAAGAAAAC
TCACAAGGCAAATATAGAGATTAGCCTTGATGAATTTAAGTTCATGTTAATGCTTGAAAATAACTACC
ATGAGTTTAAAAGGCTTAACCAATGGGTTTTGAAACCAATAAGTAAAGATTTAAACACTTACAGCAAT
ATGAAATTGGTGGTTGATAAGCGAGGCCGCCCGACTGATACGTTGATTTTCCAAGTTGAACTAGATAG
ACAAATGGATCTCGTAACCGAACTTGAGAACAACCAGATAAAAATGAATGGTGACAAAATACCAACAA
CCATTACATCAGATTCCTACCTACATAACGGACTAAGAAAAACACTACACGATGCTTTAACTGCAAAA
ATTCAGCTCACCAGTTTTGAGGCAAAATTTTTGAGTGACATGCAAAGTAAGTATGATCTCAATGGTTC
GTTCTCATGGCTCACGCAAAAACAACGAACCACACTAGAGAACATACTGGCTAAATACGGAAGGATCT
GAGGTTCTTATGGCTCTTGTATCTATCAGTGAAGCATCAAGACTAACAAACAAAAGTAGAACAACTGT
TCACCGTTACATATCAAAGGGAAAACTGTCCATATATGCACAGATGAAAACGGTGTAAAAAAGATAGA
TACATCAGAGCTTTTACGAGTTTTTGGTGCATTCAAAGCTGTTCACCATGAACAGATCGACAATGTAA
CAGATGAACAGCATGTAACACCTAATAGAACAGGTGAAACCAGTAAAACAAAGCAACTAGAACATGAA
ATTGAACACCTGAGACAACTTGTTACAGCTCAACAGTCACACATAGACAGCCTGAAACAGGCGATGCT
GCTTATCGAATCAAAGCTGCCGACAACACGGGAGCCAGTGACGCCTCCCGTGGGGAAAAAATCATGGC
AATTCTGGAAGAAATAGCGCTTTCAGCCGGCAAACCTGAAGCCGGATCTGCGATTCTGATAACAAACT
AGCAACACCAGAACAGCCCGTTTGCGGGCAGCAAAACCCGTGGGAATTAATTCCCCTGCTCGCGCAGG
CTGGGTGCCAAGCTCTCGGGTAACATCAAGGCCCGATCCTTGGAGCCCTTGCCCTCCCGCACGATGAT
CGTGCCGTGATCGAAATCCAGATCCTTGACCCGCAGTTGCAAACCCTCACTGATCCGCATGCCCGTTC
CATACAGAAGCTGGGCGAACAAACGATGCTCGCCTTCCAGAAAACCGAGGATGCGAACCACTTCATCC
GGGGTCAGCACCACCGGCAAGCGCCGCGACGGCCGAGGTCTTCCGATCTCCTGAAGCCAGGGCAGATC
CGTGCACAGCACCTTGCCGTAGAAGAACAGCAAGGCCGCCAATGCCTGACGATGCGTGGAGACCGAAA
CCTTGCGCTCGTTCGCCAGCCAGGACAGAAATGCCTCGACTTCGCTGCTGCCCAAGGTTGCCGGGTGA
CGCACACCGTGGAAACGGATGAAGGCACGAACCCAGTGGACATAAGCCTGTTCGGTTCGTAAGCTGTA
ATGCAAGTAGCGTATGCGCTCACGCAACTGGTCCAGAACCTTGACCGAACGCAGCGGTGGTAACGGCG
CAGTGGCGGTTTTCATGGCTTGTTATGACTGTTTTTTTGGGGTACAGTCTATGCCTCGGGCATCCAAG
CAGCAAGCGCGTTACGCCGTGGGTCGATGTTTGATGTTATGGAGCAGCAACGATGTTACGCAGCAGGG
CAGTCGCCCTAAAACAAAGTTAAACATCATGAGGGAAGCGGTGATCGCCGAAGTATCGACTCAACTAT
CAGAGGTAGTTGGCGTCATCGAGCGCCATCTCGAACCGACGTTGCTGGCCGTACATTTGTACGGCTCC

GCAGTGGATGGCGGCCTGAAGCCACACAGTGATATTGATTTGCTGGTTACGGTGACCGTAAGGCTTGA
TGAAACAACGCGGCGAGCTTTGATCAACGACCTTTTGGAAACTTCGGCTTCCCCTGGAGAGAGCGAGA
TTCTCCGCGCTGTAGAAGTCACCATTGTTGTGCACGACGACATCATTCCGTGGCGTTATCCAGCTAAG
CGCGAACTGCAATTTGGAGAATGGCAGCGCAATGACATTCTTGCAGGTATCTTCGAGCCAGCCACGAT
CGACATTGATCTGGCTATCTTGCTGACAAAAGCAAGAGAACATAGCGTTGCCTTGGTAGGTCCAGCGG
CGGAGGAACTCTTTGATCCGGTTCCTGAACAGGATCTATTTGAGGCGCTAAATGAAACCTTAACGCTA
TGGAACTCGCCGCCCGACTGGGCTGGCGATGAGCGAAATGTAGTGCTTACGTTGTCCCGCATTTGGTA
CAGCGCAGTAACCGGCAAAATCGCGCCGAAGGATGTCGCTGCCGACTGGGCAATGGAGCGCCTGCCGG
CCCAGTATCAGCCCGTCATACTTGAAGCTAGACAGGCTTATCTTGGACAAGAAGAAGATCGCTTGGCC
TCGCGCGCAGATCAGTTGGAAGAATTTGTCCACTACGTGAAAGGCGAGATCACCAAGGTAGTCGGCAA
ATAATGTCTAACAATTCGTTCAAGCCGACGCCGCTTCGCGGCGCGGCTTAACTCAAGCGTTAGATGCA
CTAAGCACATAATTGCTCACAGCCAAACTATCAGGTCAAGTCTGCTTTTATTATTTTTAAGCGTGCAT
AATAAGCCCTACACAAATTGGGAGATATATCATGAAAGGCTGGCTTTTTCTTGTTATCGCAATAGTTG
GCGAAGTAATCGCAACATCCGCATTAAAATCTAGCGAGGGCTTTACT
